# High rifampicin peak plasma concentrations accelerate the slow phase of bacterial decline in tuberculosis patients: evidence for heteroresistance

**DOI:** 10.1101/2022.06.06.494966

**Authors:** Antal Martinecz, Martin J. Boeree, Andreas H. Diacon, Rodney Dawson, Colin Hemez, Rob E. Aarnoutse, Pia Abel zur Wiesch

## Abstract

**Background:** Antibiotic treatments are often associated with a late slowdown in bacterial killing. This separates the killing of bacteria into at least two distinct phases: a quick phase followed by a slower phase, the latter of which is linked to treatment success. Current mechanistic explanations for the *in vitro* slowdown are either antibiotic persistence or heteroresistance. Persistence is defined as the switching back and forth between susceptible and non-susceptible states, while heteroresistance is defined as the coexistence of bacteria with heterogeneous susceptibilities. Both are also thought to cause a slowdown in the decline of bacterial populations in patients and therefore complicate and prolong antibiotic treatments. Reduced bacterial death rates over time are also observed within tuberculosis patients, yet the mechanistic reasons for this are unknown and therefore the strategies to mitigate them are also unknown.

**Methods and Findings:** We analyse a dose ranging trial for rifampicin in tuberculosis patients and show that there is a slowdown in the decline of bacteria. We show that the late phase of bacterial killing depends more on the peak drug concentrations than the total drug exposure. We compare these to pharmacokinetic-pharmacodynamic models of rifampicin heteroresistance and persistence. We find that the observation on the slow phase’s dependence on pharmacokinetic measures, specifically peak concentrations are only compatible with models of heteroresistance and incompatible with models of persistence. The quantitative agreement between heteroresistance models and observations is very good 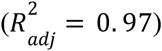.

To corroborate the importance of the slowdown, we validate our results by estimating the time to sputum culture conversion and compare the results to a different dose ranging trial.

**Conclusions:** Our findings indicate that higher doses, specifically higher peak concentrations may be used to optimize rifampicin treatments by accelerating bacterial killing in the slow phase. It adds to the growing body of literature supporting higher rifampicin doses for shortening tuberculosis treatments.

## Introduction

Tuberculosis (TB) has been afflicting human populations for millennia. The standard treatment for antibiotic susceptible tuberculosis is a 6 month long combination of rifampicin, isoniazid, pyrazinamide, and ethambutol that still carries a significant risk of relapse: up to 15% [1,2]. Shortening these treatments is a public health priority; however, this is difficult to achieve due to the lack of insight into how antibiotic therapies affect bacterial populations *in vivo* in general, and especially in the case of tuberculosis.

In clinical trials of (pulmonary) tuberculosis, the clinical course of patients and treatment efficacy are often monitored with sputum mycobacterial cultures. In phase 2A of tuberculosis drug development, i.e. early bactericidal activity (EBA) trials, quantitative sputum bacterial counts are commonly used and measured daily for up to 14 days. Later phases of drug development focus on the subsequent time period in which sputum cultures return negative and therefore the time of sputum culture conversion from positive to negative (TSCC) is used instead [3,4]. TSCC has been shown to be correlated to the probability of relapse and unfavourable outcomes [5]. This indicates that the bacterial burden in sputum may be indicative of the total bacterial burden in a patient, even if the sputum alone does not allow a complete quantification of the bacterial burden. This is because i) not all bacteria in sputum can be cultured [6–8]; ii) only bacteria in (micro)cavities connected to the airways are thought to be detected in sputum, and intracellular bacteria or bacteria in granulomas remain inaccessible; iii) even though sputum culture conversion is expected to occur in the first 2 months of treatment, patients still have to be treated for the full 6 months in order to avoid relapse.

In addition to imperfect quantification, the rate at which culturable bacteria in sputum decline is difficult to determine. This is because bacterial killing follows a typical biphasic pattern with a fast initial and a slow late decline. Such bi- or multiphasic bacterial killing has been observed both *in vivo* and *in vitro* across bacterial species. The nature and cause of these biphasic kill curves are a hotly debated subject in microbiology. The different explanations for this slowdown can be divided into two general concepts: persistence (Figure 1A) and heteroresistance (Figure 1B) [9]. The concept of persistence implicates mechanisms that rely on switching back and forth between an antibiotic susceptible and a non-susceptible state. The most commonly shown mechanisms are the switch between a replicating and a non-replicating state, and a switch between intracellular and extracellular lifestyles [10–12]. The other concept, heteroresistance, implicates mechanisms that rely on diversity in the susceptibility to antibiotics within the population of bacteria [13–15]. While in tuberculosis research, heteroresistance usually refers to the coexistence of both susceptible and resistant strains, in microbiology it also refers to the observation that not all bacteria in a clonal culture exhibit the same antibiotic susceptibility [16]. Both phenomena have been shown to exist in *Mycobacterium tuberculosis* (*Mtb*) when exposed to rifampicin *in vitro*. Both likely exist *in vivo* as well, although their clinical significance it unknown [11,17,18]. Currently, it is unclear which mechanisms contribute to the biphasic decline of bacteria, including *Mtb* in sputum. This is difficult to investigate experimentally, because of the inaccessible location of the bacteria (most often lungs, but also other inaccessible organs) and because most microbiological characterizations require removing bacteria from the patients, thereby altering both the bacteria and their microenvironment.

**Figure 1.**
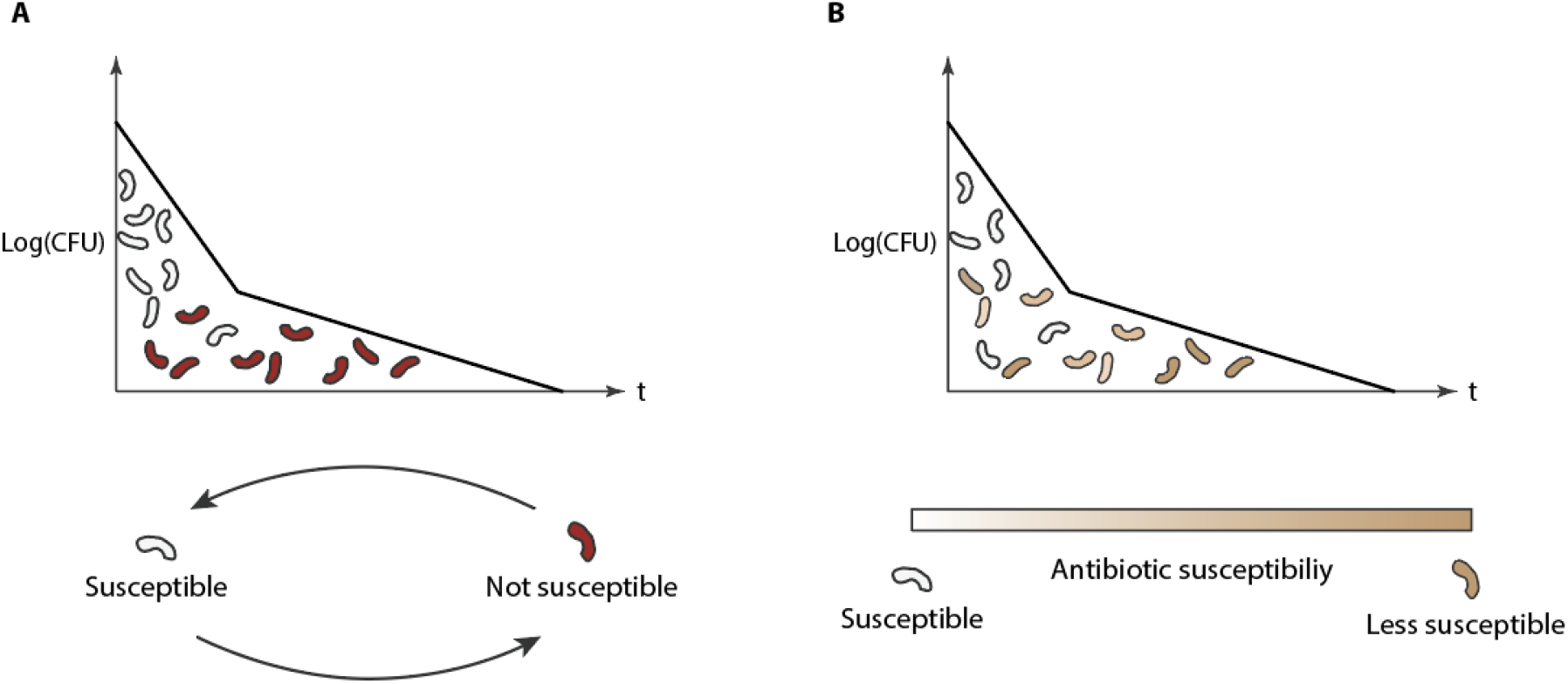
Mechanisms that can result in a slowdown in bacterial decline: persistence and heteroresistance. In both cases, X-axis is the time and the Y axis is the number of bacteria. **Figure A** illustrates persistence where bacteria switch back and forth between a susceptible and non-susceptible state over time only the non-susceptible ones remain. These are only killed when they exit a non-susceptible state. **Figure B** illustrates heteroresistance that is defined by a diversity within the susceptibility on antibiotics within the population of bacteria, some subpopulations will end up being killed at a slower pace.

Recently it was shown that the slow phase of bacterial killing may be predictive of treatment success, failure, or relapse [19]. Multiple other studies involving rifampicin, one key drug used in the treatment of tuberculosis, have linked bacterial load in sputum to treatment success. Mathematical models based on clinical trials have linked rifampicin exposure to TSCC [20], as well as to the decrease in bacterial loads measured over 14 days [21]. Another study with a limited number of patients linked higher rifampicin doses per bodyweight to the rate of late bacterial decline [13]. Animal models have also predicted a significant treatment shortening with higher rifampicin concentrations [22]. However, the results of these studies are difficult to use in mathematical models that would aid in designing optimal TB therapy.

A quantitative approach that links antibiotic exposure to bacterial killing would allow us to compare clinical results to mathematical model predictions and would thereby aid in investigating the nature of the slowdown in bacterial decline. Specifically, we would like to investigate the effects of changing antibiotic time-concentration profiles in models of persistence and heteroresistance and compare them to observed patterns in patients with different pharmacokinetic characteristics. Depending on the causes behind the biphasic behaviour, pharmacokinetic parameters may have different impact on treatment success and bacterial counts. In mathematical models of heteroresistance, we have shown previously that the slow phase can be only affected by higher drug doses if it is caused by a diversity in decline rates, rather than switching between states [13,23].

In order to uncover the mechanisms behind the slowdown in bacterial decline, we analysed the bacterial count measurements from a clinical trial on varying doses of rifampicin in tuberculosis treatment (NCT01392911, [24]). Next, we compared the results from this data analysis with mathematical models of heteroresistance and persistence. Finally, based on the statistical analysis, we aimed to estimate how TSCC may depend on treatments and compare the estimates to measured TSCCs in a different clinical trial in tuberculosis treatment (NCT01785186, [25]). Our approach allows us to show not only the possible mechanisms behind the slowdown in elimination, but also how to mitigate them.

## Materials and Methods

### Dataset

For the main analysis, we used the dataset of NCT01392911, an early bactericidal activity dose ranging trial on tuberculosis patients. Here, participants received 10, 20, 25, 30, 35 or 40 mg doses per kg bodyweight of rifampicin as a monotherapy for the initial seven days, after which isoniazid, pyrazinamide, and ethambutol were added in standard doses for the second seven days of the trial. The effects of this is discussed in the results. From this dataset, we only used the data of participants that had C_max_, AUC, as well as sputum bacterial count measurements available (n=80).

The dataset contained both time to positivity measurements as well as CFU measurements to quantify bacterial burden. Only CFU counts were used as time to positivity is challenging to convert into quantitative estimates of bacterial burden, and mathematical models are needed to convert time to positivity into bacteria per unit volume to calculate the rate of bacterial decline [26].

We estimated TSCCs (time to sputum culture conversion) using the main analysis and compared the estimates to the measured TSCCs in trial NCT01785186 [25]. This trial is a multi-arm multi-stage (MAMS) clinical trial on tuberculosis patients that included treatment arms on higher rifampicin doses. During the course of the trial, the participants received 12 weeks of experimental treatments, followed by the standard continuation phase (rifampicin and isoniazid) treatment for another 14 weeks. Here, we only used data from the control (HRZE, standard regimen n=123) and higher rifampicin dose (HR35ZE, standard regimen but with 35mg/kg rifampicin, n=63) treatment arms.

### Statistical analysis

In each patient, we had access to bacterial count determinations on 10 days. Our aim was to have at least 4 data points each for the early and late phase to estimate the rates of decline. Due to missing data, it was not possible to get reliable estimates of biphasic kill rates in substantial fraction of single patients. Therefore, we pooled bacterial counts from sputum samples from patients with similar pharmacokinetic measures. This also allowed us to include participants into the analysis that would not have enough measurements for fitting i.e. less than three days’ worth of measurements for the quick or the slow phase. We divided the range between the measured minimum and maximum pharmacokinetic measurements (either AUC or C_max_) into equal intervals. Next, using the least squares method, we fitted biphasic curves to the pooled Log (CFU) measurements from all patients within each of the intervals (see Eq (1)) for the fitted function). Finally, each of the intervals were assigned to have the median of the AUC and C_max_ values of the patients within. We used these in univariable analyses, where we tested the relationships between pharmacokinetic measures and the quick phase, slow phase, estimated last day, Log(CFU) on days 0, day 14, and day of transition from quick phase to slow phase. Here, the estimated last day as the time point where the fitted curves go below 5 CFU, which is the estimated detection limit based on [27]. For the relationships between the pharmacokinetic parameters and the quick or the slow phases we used weighted fits, where the weights were determined using the standard errors of the fits to the quick or to the slow phase 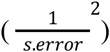. This allowed us to give smaller weights to cases where we are more uncertain about our estimates. In order to account for multiple testing, we have corrected all the P-values using the Benjamini-Hochberg method.

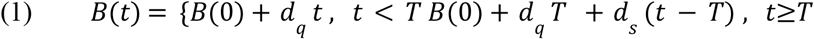

Finally, we used the corrected Akaike information criterion (AIC) values as well as the adjusted R-squared values of the fits assessing whether the C_max_ or the AUC is a better predictor for the slope of the slow phase.

### Mathematical models of the slowdown in bacterial decline

An apparent slowdown in the decline rates can stem from a diversity in the decline rates within populations of bacteria or switching back and forth between susceptible and non-susceptible states. While the two are not mutually exclusive, one or the other may dominate the response on a given timescale. In order to model this, we use mathematical models of antibiotic persistence and heteroresistance in populations of bacteria in order to be able to assess the response of the two different mechanisms to different pharmacokinetic measurements. We are using these as a reference as both of these mechanisms has been observed in mycobacteria previously [15,28]. Table 1 summarizes the parameters and values used in the equations.

**Table 1.**
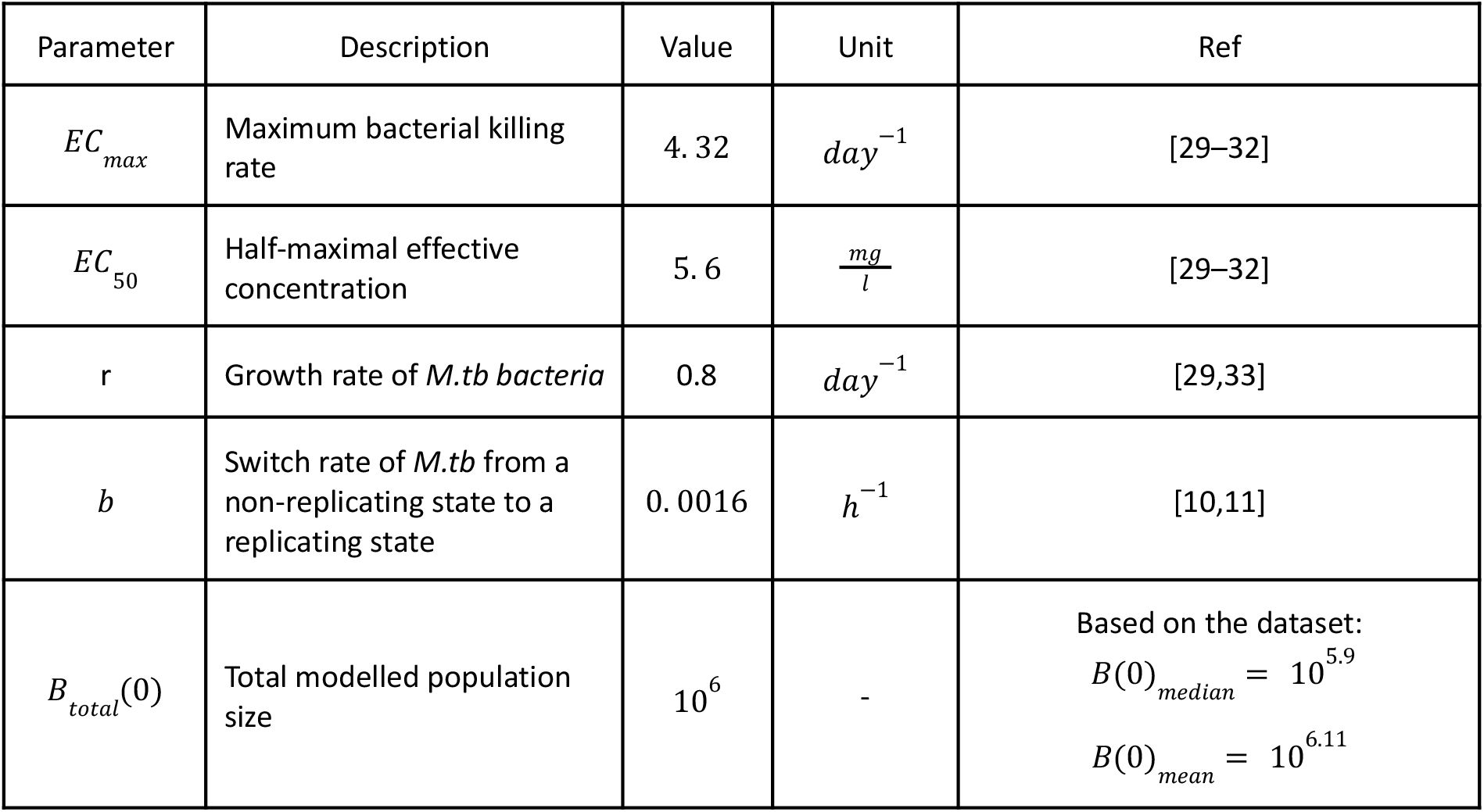
Summary of parameters used in the mathematical models of persistence and heteroresistance.

| Parameter | Description | Value | Unit | Ref |
| --- | --- | --- | --- | --- |
| $EC_{max}$ | Maximum bacterial killing rate | 4.32 | $day^{-1}$ | [29–32] |
| $EC_{50}$ | Half-maximal effective concentration | 5.6 | $\frac{mg}{l}$ | [29–32] |
| $r$ | Growth rate of *M.tb* bacteria | 0.8 | $day^{-1}$ | [29,33] |
| $b$ | Switch rate of *M.tb* from a non-replicating state to a replicating state | 0.0016 | $h^{-1}$ | [10,11] |
| $B_{total}(0)$ | Total modelled population size | $10^6$ | - | Based on the dataset:<br>$B(0)_{median} = 10^{5.9}$<br>$B(0)_{mean} = 10^{6.11}$ |

### Elimination rate curves

For both of the models, we used the antibiotic concentration – net growth (decline) rate relationship available in the literature in [32] that processed data of *in vitro* M.tb exposure to rifampicin measurements by [30,31]:

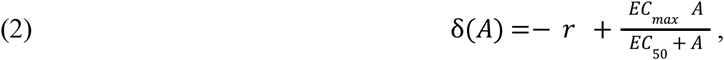

where *A* is the antibiotic concentration in 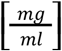.

### Phenotypic switch

The most well-known example of switching back and forth between states is bacterial persistence caused by a dormant state [10]. Here, it is assumed that a subset of bacteria is in a non-replicating, dormant state that protects them from the effects of antibiotics that damage growing bacteria (for example rifampicin). Bacteria in their dormant state should be completely or highly tolerant to antibiotics, however once they switch back to a replicating state, they are swiftly killed by antibiotics. Mathematically this can be modelled as described in Eq (3) [10]. These equations describe the two subpopulations of bacteria, one that is susceptible (*n*) and one that is non-susceptible (*p*). Once the dormant state bacteria start replicating again, they become susceptible to antibiotics again (switching rate *b* in the equations).

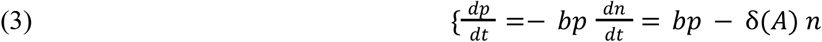

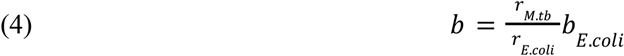

In [10], they have determined the switch rate (*b*) from non-replicating state to a replicating state for *E*.*coli* (persisters) to be. We have scaled this value by the differences in replication rates between *E*.*coli* and *M*.*tb* (Eq 4)) in order to preserve the assumptions of the original model on the determined fraction of persisters. This is supported by the literature [11] that found > 20 *day* lag times for *M*.*tb* (compared to our estimate of 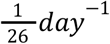 switch rates)

We chose to model the non-replicating or slowly replicating *M*.*tb* as non-susceptible to antibiotics, based on the current definition of persistence [9]. This is also supported by the literature that reported 17x, 50x, and 200x decrease in susceptibility to rifampicin in a non- or slowly replicating state [18,34,35]. This makes the decline rates of persisters negligibly small on the MIC ranges (1 – 16x MIC) we use the mathematical model on.

### Heteroresistance

The other mechanism responsible for the slowdown in the decline is heteroresistance [9,17]. The underlying assumption is that the bacteria in a population are not completely identical or may express unstable resistance genes that can result in a diversity in the susceptibility to external antibiotic concentrations. Additionally, there can also be a diversity in the expression of efflux pumps, targets, or cell sizes for example [13–15] which can be described by the same model.

Mathematically, this can be calculated using Eq (5) [13,23] that describe multiple subsets of bacteria each with a slightly different susceptibility to antibiotics.

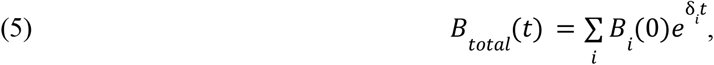

here *Bi*(0) represents the initial size of the *i*-th subpopulation. Furthermore, *δ* _*i*_ are the corresponding decline rates. The model simulates 9 subpopulations where the middle one (5^th^) has the “average” susceptibility and in the majority.

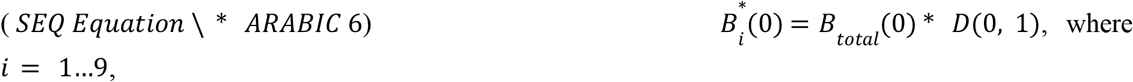

where *D(x* |*μ,σ)*, is the probability density function of the normal distribution, with the mean: μ and standard deviation σ of the distribution.

Due to the way this is calculated (by taking the value of the normal distribution at the given point), 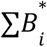 will be less than *B*_*total*_ and therefore has to be rescaled:

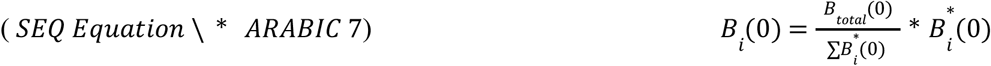

Here we simulate decrease susceptibility as changing the effective antibiotic concentrations (*A*_*i*_). For the purposes of this paper the two are the same, but this approach is easier to compute [23].

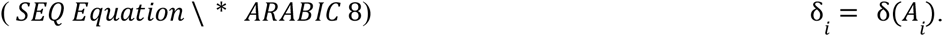

Therefore we use the following to simulate 9 subpopulations with MICs ranging from 1/16^th^,1/8^th^… to 8x and 16x of the average:

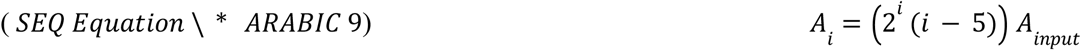

For simulations, the parameters for the population distribution *D*(*x* μ σ) are the fits to the statistical analysis as shown in Figure 4 (μ = 0, σ =1.3).

### Determining the day of transition from quick phase to slow phase

*In vivo* data is noisy, and the transition from quick phase to slow phase (based on mathematical models) is smooth rather than a sudden. Therefore, it is difficult to find the day of transition based on individual fits of the biphasic curves. We have opted for determining the day of transition based on all the available fits rather than optimizing fits in for each biphasic curve individually. Therefore we repeated the fitting procedure for all dosing groups with a different day of transition. We chose the day of transition where median R^2^ for the fits of the slow and fast decline were highest. We got the same result when minimizing the P-values as well.

### PK-PD models

The pharmacokinetic model used the published compartmental pharmacokinetic model of [36]. For the sake of simplicity we only used absorption (Eq (10)), plasma (Eq (11), and tissue (Eq (12)) compartments and have not used a compartment chain for the absorption:

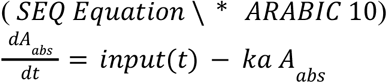

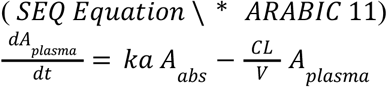

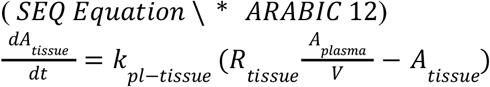

Here the parameters are:

- *input*(*t*) is the input function, to simulate daily doses of rifampicin. It implemented as Dirac delta functions (Dirac comb) spaced 24h apart.
- *A*_*abs*_,*A*_*tissue*_, and *A*_*plasma*_ are antibiotic concentrations in the absorption, plasma and tissue compartments
- *ka* = 1. 55 [*h*^−1^]: absorption rates of the drug from absorption compartment
- 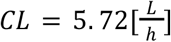 are clearance rates of the drug from the plasma compartment
- *V* = 52. 3 [*L*] is the volume of distribution in litres
- *R*_*tissue*_ is the penetration coefficient into the given tissue, see Table 2.

**Table 2.** This table details R_tissue_ and k_pl−tissue_ parameters for used throughout this work [36]
*k*_*pl*-*tissue*_ rate of drug moving from plasma to tissue (*h*^-1^), see Table 2.

| Parameter | Lung | Necrotic nodule | Caseum closed nodule | Caseous fibrotic nodule | Caseum from cavity | Cavity wall | Fibrotic tissue | Small cellular nodule | Fungal ball |
| --- | --- | --- | --- | --- | --- | --- | --- | --- | --- |
| R | 0.69 | 0.569 | 0.443 | 0.733 | 0.449 | 0.614 | 0.539 | 0.348 | 0.447 |
| k | 1.68 | 0.751 | 0.367 | 0.367 | 0.204 | 1.98 | 0.209 | 4.67 | 0.212 |

For the pharmacodynamic models, we omitted bacterial replication from the mathematical model for the sake of simplicity as well as the lack of data on the progeny of the less susceptible subpopulations as well as the persistent bacteria. Arguably there is very little replication taking place over the modelled time period (14 days). This is due to antibiotic concentrations being close to or above MIC for the majority of time, post-antibiotic effect (lag time after antibiotic exposure before the replication restarts), and the slowness of replication in vivo environments. Here, the bacterial population sizes are calculated with Eq (13):

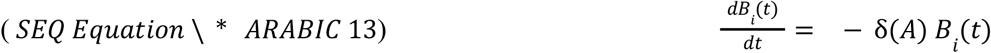

## Results

### Statistical analysis of the dataset

In this work we investigated possible mechanistic causes underlying the observed slowdown of bacterial decline rates in patients treated with different doses of rifampicin. We investigated the properties of the slowdown in elimination in response to different pharmacokinetic measures (total exposure in plasma or AUC; and peak plasma concentrations or Cmax) via statistical analysis, as different behaviours hint at the mechanistic causes of the slowdown. Here, a per-patient analysis is not viable, as fitting a distinct quick and slow phase to 10 days’ worth of data (4-6 points per phase) is unreliable due to the inherent noise of *in vivo* measurements. To resolve this, we pooled data together from multiple participants. In order to avoid bias from the choice of group sizes, we repeated the analysis with 30 different group sizes. We formed these groups by dividing the measured range of pharmacokinetic parameters (either Cmax or AUC) into 10 to 40 equal sized intervals. This ensured that a detected slowdown in elimination is consistent across patients with similar pharmacokinetic measurements, and the groupings do not introduce bias into the analysis. Unless otherwise indicated, we report the median values for all of the statistical descriptors (e.g. R^2^, p-value), and estimated parameters (reported in Table 3).

**Table 3.** Summary of the median fitted values to the dataset across all groupings, with day 4 as the day of transition from quick phase to slaw phase. All the P values reported are corrected for multiple testing with the Benjamini-Hochberg method. Here the overall decline rates (used in Eq (1)) within each phase can be calculated as: 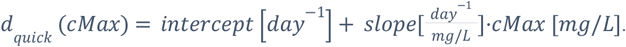

| Property | Grouped by | Intercept (SE) | Slope (SE) | Intercept p-value (corrected for multiple testing) | Slope p-value (corrected for multiple testing) | R<sup>2</sup> (adjusted) | AIC (corrected) |
| --- | --- | --- | --- | --- | --- | --- | --- |
| Quick [day<sup>-1</sup>] | AUC | -0.19 (±0.038) | -0.0005 (±0.00009) | 0.0003 | 0.00007 | 0.67 | -25 |
|  | C<sub>max</sub> | -0.1 (±0.057) | -0.0057 (±0.001) | 0.1 | 0.00016 | 0.54 | -22 |
| Slow [day<sup>-1</sup>] | AUC | -0.075 (±0.024) | -0.00015 (±0.00005) | 0.007 | 0.01 | 0.32 | -43 |
|  | C<sub>max</sub> | -0.043 (±0.026) | -0.0018 (±0.00051) | 0.12 | 0.003 | 0.34 | -57 |
| Baseline<br>Log(CFU)<br>[ - ] | AUC | 5.54<br>($\pm 0.18$) | 0.0012<br>($\pm 0.00044$) | $10^{-15}$ | 0.0124 | 0.30 | 23 |
| | C<sub>max</sub> | 5.27<br>($\pm 0.254$) | 0.016<br>($\pm 0.0052$) | $10^{-13}$ | 0.0107 | 0.27 | 40 |
| 4<sup>th</sup> day<br>Log(CFU)<br>[ - ] | AUC | 4.77<br>($\pm 0.26$) | -0.0009<br>($\pm 0.00063$) | $10^{-11}$ | 0.163 | 0.30 | 23 |
| | C<sub>max</sub> | 4.86<br>($\pm 0.307$) | -0.008<br>($\pm 0.0064$) | $10^{-12}$ | 0.23 | 0.27 | 40 |

For each groups’ pooled sputum bacterial count measurements (for all groupings), we fitted biphasic curves with different days of transition from the quick killing to the slow killing of the bacteria. We determined the most probable days of transition, day 3 or 4, based on the median R^2^ (see Figure S 1) and median p-values of the fits (see Figure S 4). For the rest of the analyses, we used day 4 as the day of transition between the quick and the slow phase.

Based on the fitted curves, we found that both phases of decline are dependent on antibiotic concentrations and therefore pharmacokinetic measures. This is demonstrated in Figure 2 which illustrates the dependence of kill rates on AUC or Cmax The resulting smoothed time-kill curves based on the fits are shown in Figure 2E. Table 3 summarizes the properties of the fitted curves, Figure S 3 shows all the determined fits for all different transition days, and Figure S 4 shows the corresponding p-values. We applied the Benjamini-Hochberg correction to account for multiple hypothesis testing when calculating all p-values reported in Table 3 and Figure S 4.

**Figure 2.**
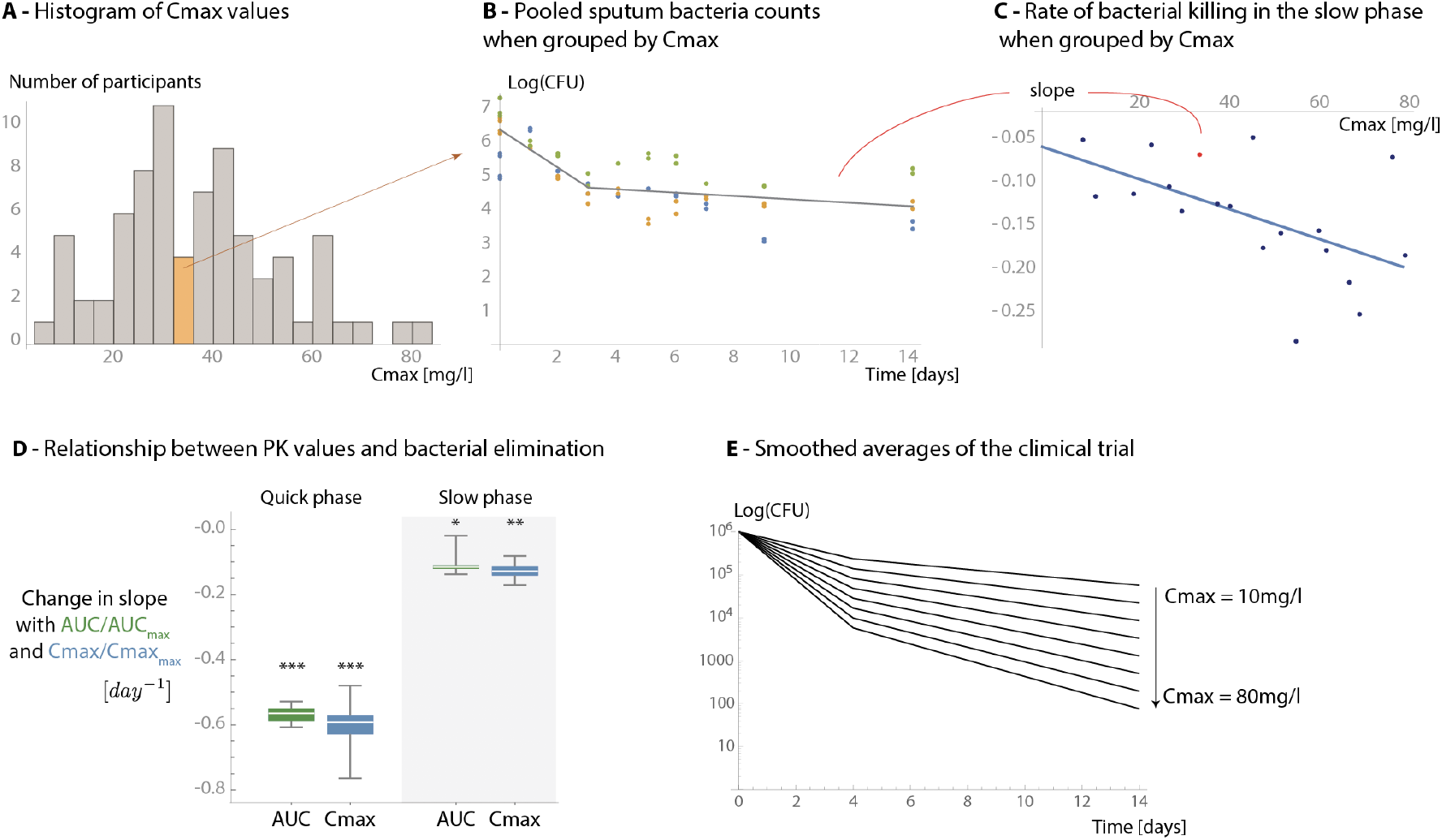
Overview of the fitting process and the relationship between the decline rates of bacteria and pharmacokinetic measures. We have grouped participants (**Figure A**) by forming equal groups within the pharmacokinetic measures. For each group, we pooled the bacterial count measurements (Y-axis) of all participants (dots, here one colour corresponds to one participant) and fitted biphasic curves to them (solid line) (**Figure B**). Next, we analysed the relationship between the groups’ median pharmacokinetic measurements (X axis) and the properties of the fitted curves (Y axis) (**Figure C**). **Figure D** summarizes the dependence of the decline rates in the quick and slow phases on 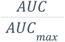 and 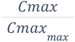. Here, we have normalized the values (by dividing with the maximum), in order to be able to show that both measures produce the same result but with different errors. **Figure E** demonstrates how this affects the bacterial count measurements (X-axis) over time (Y-axis). The curves were made with Eq (1), starting at 10^6^ CFU/ml bacteria, using day 4 as the day of transition, with the obtained fits for decline rates (see Table 1), and Cmax values of 10, 20, 30…80 [mg/l] (in the dataset the Cmax values range from 7.7 to 85.6 mg/ml).

### Mathematical modelling of heteroresistance and persistence

We compared the predictive power of the pharmacokinetic measures using the adjusted R2 (Figure S 1) and corrected Akaike Information Criterion (AIC) values (Figure S 2) of the fits. The quick phase was better predicted by the AUC rather than the Cmax, while the slow phase was better predicted by the Cmax rather than the AUC. The dependence on pharmacokinetic measures of the quick phase was expected as rifampicin efficacy is thought to be better predicted by the AUC [37]. However, the slow phase’s dependence on Cmax was previously unknown and can both aid in optimizing therapy and give insight into the possible mechanistic causes of the slowdown. We demonstrate this in Figure 3 in which we modelled the responses of the two main possible causes of the slowdown (persistence and heteroresistance) when *M. tuberculosis* is exposed to rifampicin. We used simplified exposure profiles in which the AUC remains constant: a sustained high concentration (Cmax) for shorter duration of time and a low sustained concentration for longer duration of time. We found that heteroresistance predicts that antibiotic concentrations affect the slope of the slow phase, while persistence predicts that antibiotic concentrations will have no effect on the slope of the slow phase. The simplified cases also demonstrate that a high Cmax is required to kill the less susceptible subpopulations in heteroresistance. Therefore, our observations on the properties of the slow phase are consistent with the definitions and models of heteroresistance and are inconsistent with the definition and mathematical models of persistence [9].

**Figure 3.**
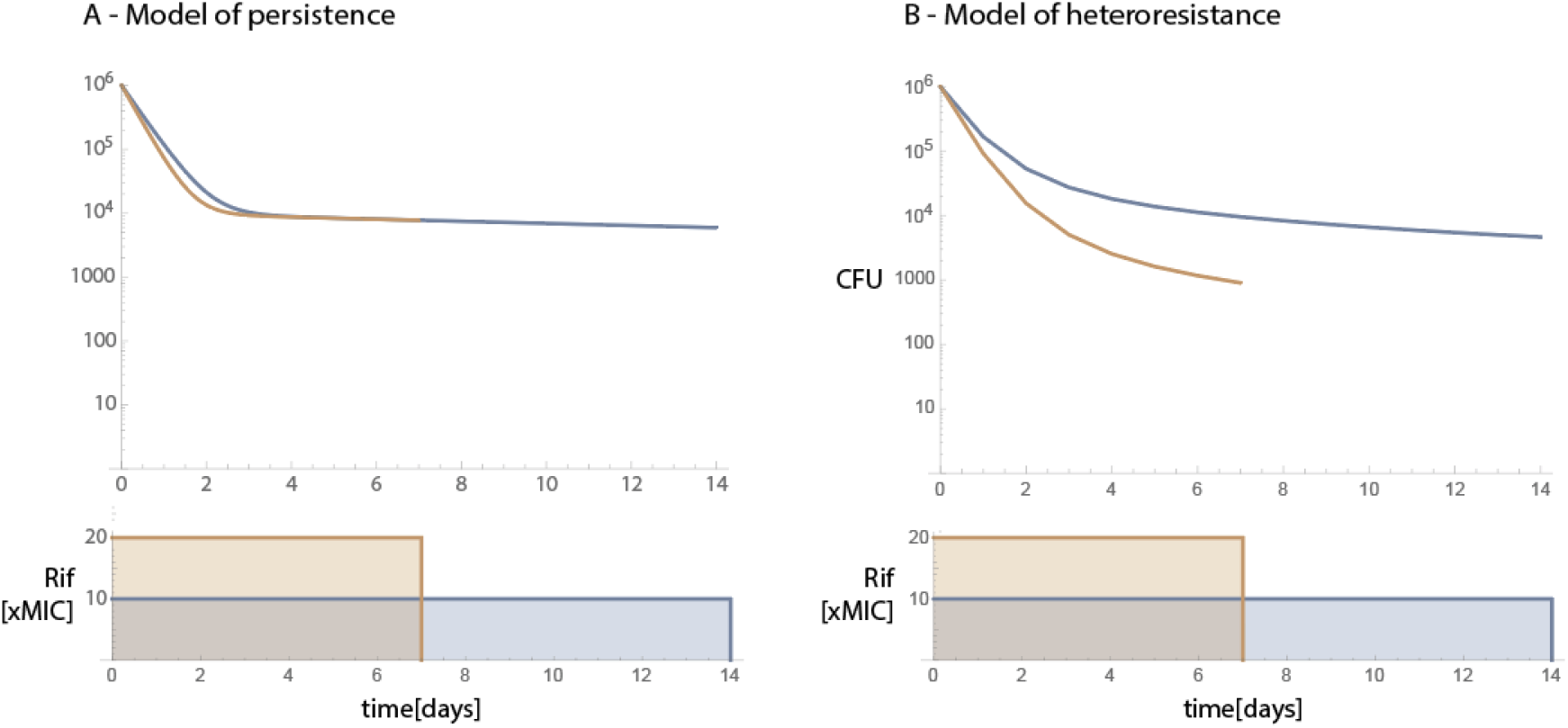
Predicted responses of the mathematical models of persistence and heteroresistance. This figure shows the qualitative difference between the causes behind the slowdown in bacterial decline. The top row shows bacterial counts (Y-axis) over time (X-axis) for two different idealized dosing regimens (grey and black curves). The bottom row shows the two simplified exposure profiles: both have the same AUC but different Cmax values. **Figure A** shows that if the slowdown in decline is a result of switching between susceptible and non-susceptible states, then the slope of the slow phase is not dependent on C_max_. On the other hand, **Figure B** shows that if the slowdown in decline is the result of a diversity in the susceptibility to antibiotics, then a change in antibiotic concentrations should affect the slope of the slow phase as well.

Heteroresistance is defined as the coexistence of multiple subpopulations with different susceptibilities, where some (minority) subpopulations have at least 4 or 8x the MIC when compared to the majority of the population [9,38]. Therefore, to further assess whether the slowdown in decline is caused by heteroresistance we calculated the hypothetical sensitivity distribution of bacteria based on the observed slowdown. Figure 4A and B illustrate the approach, which is based on combining PK-PD modelling with the results of the statistical analysis of the clinical trial. We calculated how subpopulations with different sensitivities in cavity walls would be killed with daily doses of rifampicin and compared this to the decline rates of the slow phase in the clinical trial at each level of Cmax. This allowed us to derive the sensitivity of the subpopulation that dominates the slow phase. We assigned the corresponding subpopulation size (based on the dataset Figure 4C) to each subpopulation that yielded the sensitivity distribution of bacteria (Figure 4D, purple dots). The resulting distribution shows a strong agreement with the definitions of heteroresistance (4-8x MIC subpopulations). In persistence, the size of the subpopulation with a slow decline should be independent of the antibiotic concentrations (i.e. Figure 4C would show different initial subpopulation sizes and the late bacterial decline rates would be indistinguishable).

**Figure 4.**
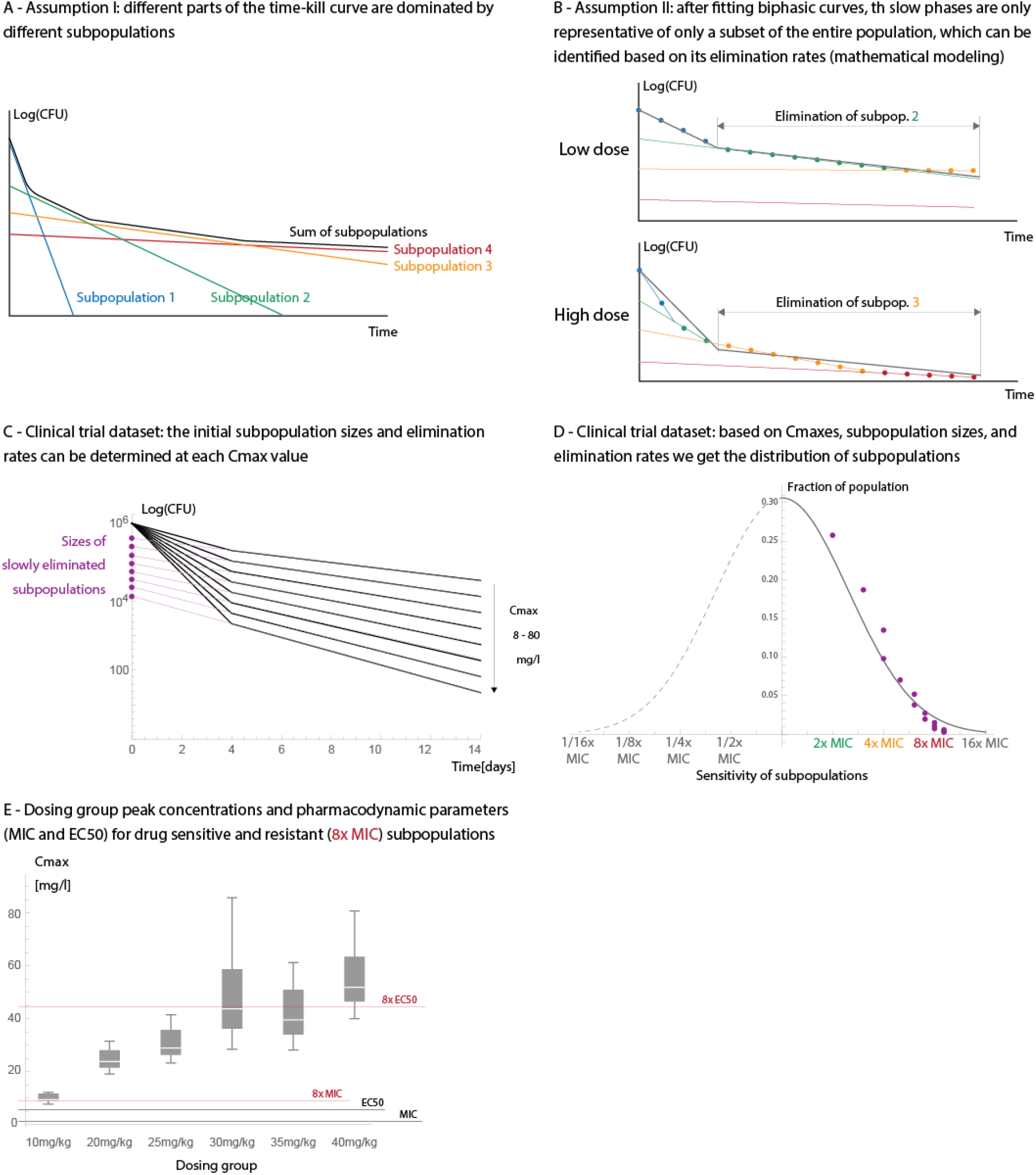
Estimating the sensitivity distribution of bacteria. This figure shows the method we used to estimate the sensitivity distribution of bacteria when the slowdown is caused by heteroresistance. **Figure A (illustration)** shows that different parts of the time-kill curves (bacterial count (Y-axis) over time (X-axis)) represent the decline of subpopulations with different MICs (different colours) [23]. In these cases, even after fitting biphasic curves on a multi-phasic time-kill curves (**Figure B (illustration))**, the slow phase will represent a subset of subpopulations which can be identified based on the decline rates at the given antibiotic concentration (Cmax or AUC in case of PK based models). **Figure C** shows the parameters used obtained from clinical trial dataset: decline rates in the slow phase and the corresponding “subpopulation sizes” at each Cmax. **Figure D** shows the obtained sensitivity distribution and the fitted (assumed) normal distribution to it. **Figure E** shows the distribution of Cmax values in each dosing group and how they compare to the pharmacodynamic parameters of MIC (1.3mg/l) and EC50 (5.6mg/l, concentration at which bactericidal activity is 50%) [32]. These values are marked for both a drug sensitive subpopulations and a less susceptible subpopulation (8x MIC) to illustrate how the sensitivity of bacterial subpopulations in Figure D relate to the pharmacodynamic parameters. Here, we assume that the EC50 values of the less susceptible (8x MIC) subpopulations are 8x the original EC50 values. However, the correct relationship between MIC and EC50 in different subpopulations is unknown and this is for illustrative purposes.

We also compared how well the PK-PD model of heteroresistance predicts the observed slowdown in decline when compared to the smoothed averages of the clinical trial dataset. To do so, we used the same model as for determining the sensitivity distributions. The parameters of the mathematical model are based on the literature: the pharmacokinetic model is based on [36] and the pharmacodynamic parameters on [32]. Figure 4 shows the sensitivity distribution in heteroresistance, which is fitted to the clinical trial dataset (assumed Gaussian distribution σ truncated at 16x MIC, see black line Figure 4D). Figure 5A, B, and C show side by side the decline rates of bacteria in the clinical trial, as well as the predicted decline rates by heteroresistance and for persistence. Figure 5D demonstrates that when compared, the difference between the observed decline of bacteria in the clinical trial and the one predicted by heteroresistance stay within one order of magnitude (see Figure S 5 for individual comparison of the curves). Therefore, we conclude that this model can sufficiently explain the observed slowdown on this timescale.

**Figure 5.**
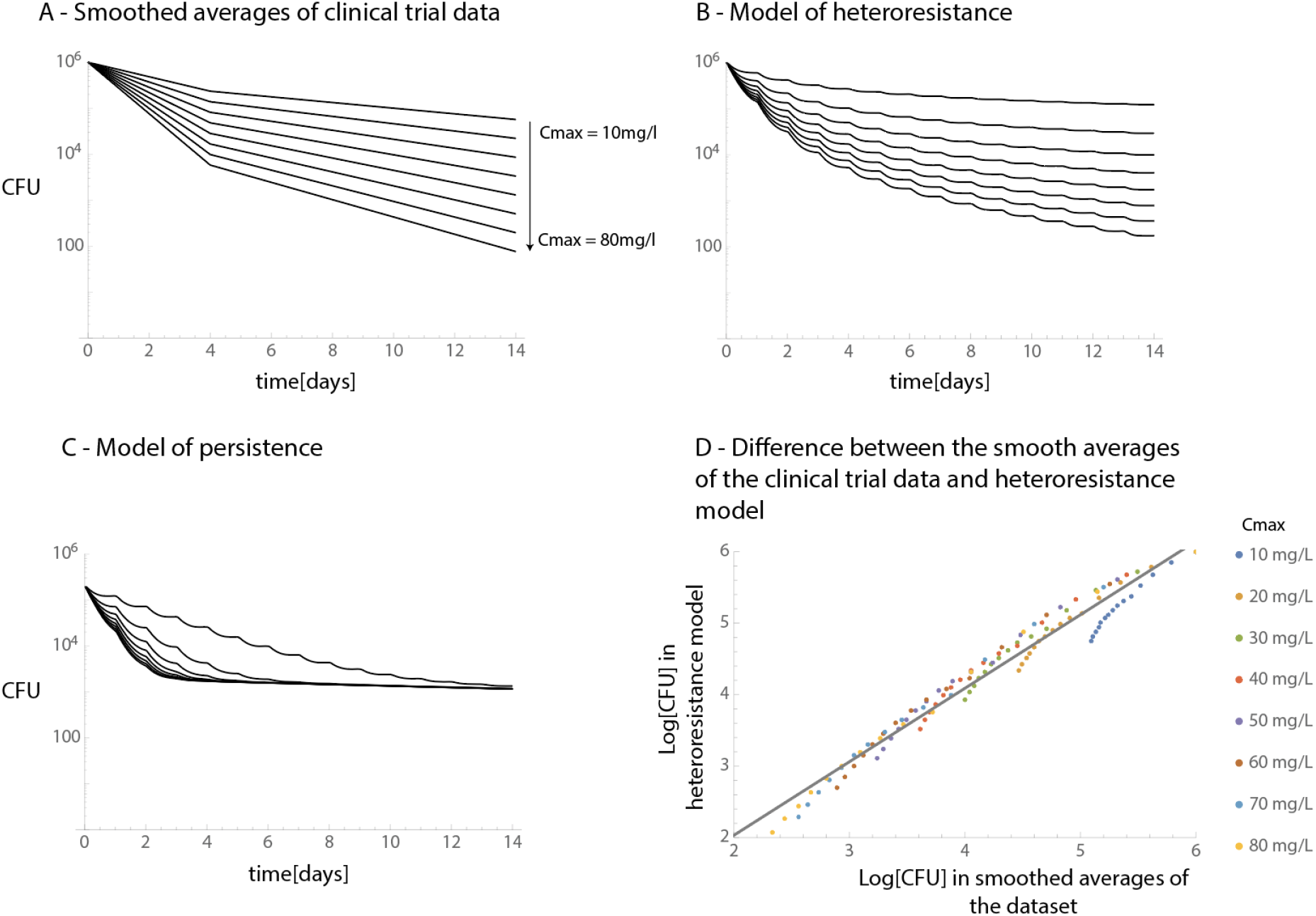
Comparison of the smoothed averages of the clinical trial and model predictions on sputum bacterial counts over time. This figure compares the smoothed averages of the clinical trial (**Figure A**, this is the same plot as in Figure 2E), the predicted decline of bacteria in cavity walls if the slowdown is caused by heteroresistance (**Figure B**) or persistence (**Figure C**). We generated these plots using pharmacokinetic models in the cavity wall with the same Cmax values as in Figure A (10…80 mg/l). Finally, **Figure D** shows the difference between the curves on Figure A and B: on all days and Cmax values combined, the X axis shows the model predictions and the Y axis shows the smoothed averages of the clinical trial 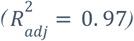.

In the PK models, Cmax and AUC both depend on multiple parameters and are only mildly correlated to each other. An increase in AUC may lower the Cmax. As a result, when the Cmax values are close to the MIC the drug concentrations might not be high enough to overcome the MIC and the bactericidal action can decrease dramatically. Figure 4E demonstrates this, there we plotted the relationship between the observed peak drug concentrations in the clinical trials and the pharmacodynamic parameters of MIC and EC50. The latter, EC50 is the concentration at which the elimination rate of bacteria is 50% of the maximum. This shows that for the 10mg/kg group, even in the plasma, the 8x MIC may not be reached. Concentrations at which bacterial elimination of a less susceptible subpopulation is “efficiently eliminated (i.e. at the peaks there is briefly 50% the maximum elimination rate)” may not be reached in the dosing groups below 30mg/kg.

### Estimating the time to sputum culture conversion

Finally, we estimated the expected time to sputum culture conversion based on the statistical analysis, assuming that there is no additional slowdown outside the timeframe of the clinical trial. We defined this as the timepoint where the fitted curves would go below 5 CFU, based on the estimated detection limits described in [27]. Figure 6 shows the expected time to culture conversion depending on Cmax and AUC. We repeated the statistical analysis using monophasic kill curves (i.e. neglecting the possibility of a slowdown in bacterial decline (see Table S1) and estimated TSCC that way as well. We compared both estimates to the measured times to sputum culture conversion of a multi-arm multi-stage (MAMS) clinical trial on higher rifampicin doses (NCT01785186, [25]). From the MAMS trial, we used the higher rifampicin (HR35ZE, standard treatment with an increased, 35mg/kg rifampicin dose) arm. Estimates that take the slowdown in bacterial decline into account (biphasic rather than monophasic fits) provide estimates that are closer to the observed data and are more conservative. In addition we also compared estimates based on the standard dosing (HRZE, standard treatment with 10mg/kg rifampicin) treatment arm. There, the Cmax and AUC range of the MAMS trials were outside the range of the Cmax and AUC values of the EBA trial we fitted the model to. Therefore, these comparisons are provided for completeness’ sake and can be found in the supplement (Figure S6). For the HR35ZE treatment arms, predictions of TSCC do not differ significantly between Cmax and AUC as in both cases concentrations are well above the MICs for most subpopulations in most of the anatomical sites and therefore are expected to be similar.

**Figure 6.**
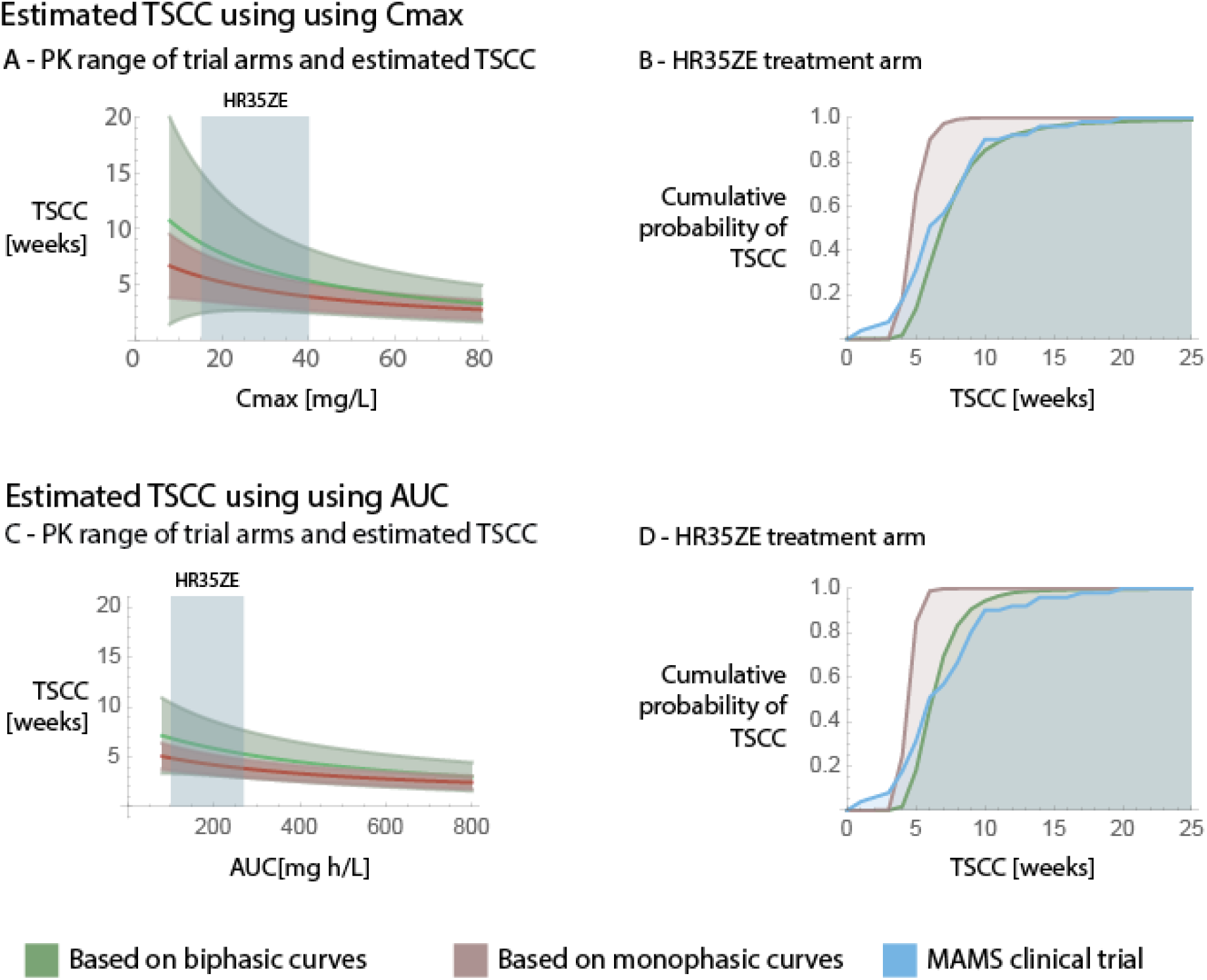
Estimated TSCC based on the statistical analysis, and comparison to measured TSCCs in a multi arm multi stage (MAMS) clinical trial (NCT01785186, [25]). Here, the green curves show estimates based on biphasic curves (i.e taking the slowdown in decline into account), while red shows estimation based on monophasic curves (i.e. neglecting the possibility of a slowdown in decline). Blue always corresponds to data from the MAMS clinical trial. **Figures A** and **C** show the dependence of the predicted TSCC on the pharmacokinetic parameters (Cmax and AUC respectively), as well as the measured pharmacokinetic ranges for the HR35ZE treatment arm of the MAMS clinical trial used for comparison (blue boxes). Here, the area around the estimates signify the 95% confidence interval around our estimates. The estimates are only shown within the parameter ranges of the EBA clinical trial. **Figures B and D** show the cumulative probability (Y-axis) of TSCC (X-axis) based on the estimates within the PK ranges of the MAMS trial as well as the data from the MAMS trial itself.

## Discussion

We analysed the measured sputum colony counts in a rifampicin dose ranging measuring early bactericidal activity (EBA) and found that there is a slowdown in the bacterial killing rates, resulting in distinct quick (days 1-4) and slow phases (days 5-14). These phases correlated significantly with patient rifampicin plasma concentrations. In case of the slow phase, this was previously unclear as the dependence on pharmacokinetic parameters of the slow phase can depend on the mechanism causing the slowdown. Therefore, it may or may not respond to increased antibiotic concentrations. Furthermore, we found that in the slow phase, the peak drug concentration (Cmax) is a better predictor for bacterial killing than the overall exposure (AUC). We have done this by showing a direct statistical link between measured pharmacokinetic parameters and sputum colony counts at different stages of decline.

This analysis of the dataset can be used to differentiate between the mechanistic causes of the slowdown. In microbiology, there are two main explanations for a slowdown in decline of bacteria: antibiotic persistence and heteroresistance [9]. Both persistence and heteroresistance have been consistently shown to exist even in cultures of clonal bacterial populations, including *M*.*tb* [11,15,28,39]. Additionally, both are not mutually exclusive, can act on different timescales and are dependent on the mode of actions of the antibiotics used. While the clinical significances of both phenomena are unknown [9,38,40], both have been shown to exist in in tuberculosis patients [11,17,28,41–43]. Most recently, differences in rifampicin response in tb patients was linked to heterogeneity in the bacterial population using NGS, however the mechanism (heteroresistance or persistence) was not described [44]. In persistence, the slow phase is driven by bacteria exiting a non-susceptible state which is commonly assumed to be independent of antibiotic concentrations. Therefore, higher antibiotic concentrations would not accelerate the slow phase of decline to the extent observed in the dataset [9]. In heteroresistance, the slow phase is driven by subpopulations that are less susceptible to antibiotics than the majority of the bacterial population. Therefore, higher antibiotic concentrations would accelerate the slow phase decline similarly to the dependence observed in the dataset [9,13,23,38]. The findings from the data analysis are corroborated by the concentration dependences observed by others as well [27]. Our statistical analysis has shown that the results are only compatible with the definitions of heteroresistance and not persistence.

To validate the findings based on the analysis of the dataset we compared the results to mathematical models of both persistence and heteroresistance parametrized with data available in the literature. The pharmacodynamics component is based on *in vitro* data of *M*.*tb* exposed to rifampicin [29–32]. The pharmacokinetic model (where applicable) is based on models of rifampicin tissue concentrations in TB patients [36]. For the simulations we used cavity walls as open cavities are thought to be one of the sources of sputum. We found that the most probable cause of the slowdown in decline on this time scale (in response to rifampicin) is heteroresistance rather than antibiotic persistence. An alternative hypothesis to this is that the slowdown in bacterial decline is caused by the heterogenous distribution of drugs in the lungs that results in diverse decline rates in bacteria [41,45,46]. In our case, this scenario is unlikely, as our results indicate that some subpopulations are likely to have 8x the MIC meaning that some tissues would have 1/8^th^ of the drug concentration. The variation in exposure levels in different tissues cannot explain this large difference [36]. Additionally, the different environments in different anatomical sites could result in different susceptibilities to antibiotics as well [41], however there are no available pharmacodynamic models on this that could be used for modelling. As this is a possibility, a more cautious summary of our results is that the slowdown in elimination is caused by heterogeneous bacterial elimination rates within the compartments from which the sputum is originating.

Through mathematical modelling we also show that the observed PK dependence of the quick and the slow phases are consistent with heteroresistance. In heteroresistance, the quick phase represents the elimination of the majority of the bacterial populations, where of the decline should be better predicted by the AUC than Cmax due to the AUC dependence of rifampicin [37]. In the slow phase however, the Cmax should be a better predictor than the AUC (Figure S 2). This is because the slow phase represents the decline of the less susceptible subpopulations where an increased MIC has to be overcome. Increasing AUC can potentially decrease Cmax which in turn may reduce the ability to overcome the higher MICs of some subpopulations. This is also what we have found in the dataset where the quick phase showed a stronger relationship with the AUC while the slow phase showed a stronger relationship with the Cmax. These differences were observable despite the fact that only one drug formulation and dosing regimen was used in the clinical trial and therefore the Cmax and AUC measurements are highly correlated. As a consequence, the analysis on the differences between the effects of Cmax and AUC rely on the patient-to-patient variance in the drug absorption, elimination, and distribution rates. We expect the differences between Cmax and AUC to be more pronounced in trials that vary drug formulations or dosing strategies as well as drug doses. This highlights the possibility of further optimizing treatments and are in support of high rifampicin peak concentrations. The current higher dose rifapentine trials (NCT02410772 [47]) can provide more clues on optimization once they are published as rifapentine has a different pharmacokinetic profile than rifampicin. However, it is difficult to predict which type of time-concentration profile would be ideal. First, it is unclear whether the patient’s tolerability depends on AUC or Cmax. Second, it has been argued that antibiotics used in tuberculosis treatments should be matched with others that have similar half-lives to avoid only one drug being present (functional-monotherapy [36]) and thereby not facilitating the emergence of resistance.

The clinical significance of heteroresistance is still unknown [38,40], however we demonstrate that a slowdown in bacterial decline due to heteroresistance delays the time to sputum culture conversion (TSCC) and therefore may reduce treatment success rates. An earlier TSCC is mildly associated with treatment success and the lack of it is associated with unfavourable outcomes and are important biomarkers in tuberculosis research [5,48]. We estimated TSCC based on the statistical analysis that assumes that there is a slowdown in bacterial decline, then we repeated the analyses by assuming that there is no slowdown (i.e. fitting a straight line to the bacterial counts) as a control. Next, we compared these estimates to the measured TSCCs from a multi arm multi stage (MAMS) clinical trial on high rifampicin doses. We show that with the use of biphasic curves the estimates on TSCC are not only more conservative but show better agreement with the data used for comparison. This validates our analysis and shows that the slowdown in elimination is present during treatments. It also serves as a demonstration on the impact of neglecting the possibility of slowdown on assessing experimental treatments. In concordance with the recently published paper using data from the first 8 weeks of treatment, our observations based on the first two weeks imply that the decline rate may not change significantly between 2-8 weeks. These results argue against the practice of having more sparse bacteriological assessments in the second half EBA clinical trials (as it was done in this trial as well), because it makes investigating the possible slowdown in bacterial decline more difficult.

These findings strengthen the connection between quantitative bacteriological assessments in EBA trials (sputum bacterial count measurements) and the TSCC that is used in subsequent phases of trials. The importance of strengthening this connection has been highlighted in the recent literature as one of the current challenges in tuberculosis clinical trial design [3,4]. We also identify a possible source of errors when evaluating EBA trials: not all drugs or drug combinations may result in the same slowdown in bacterial killing. Additionally, a slowdown may be outside of the timeframe of the two weeks of the trial (for example in bedaquiline [49]), causing us to significantly underestimate TSCC. Conversely, some drug combinations may cause little to no slowdown – in EBA trials, these may appear inferior to other treatment arms that have a fast initial decline of bacteria and a slowdown later on.

This work has several limitations as well: first, due to the number of participants in the study it was not possible to correct for the observed association between the baseline bacterial loads and the PK values. While it does not affect the calculation of decline rates itself, it may be a confounder [50,51]. Second, in the EBA trial during the first week, all the participants received rifampicin monotherapy; in the second week they have also received the standard doses of the remaining TB drugs (isoniazid, pyrazinamide, ethambutol) in combination with rifampicin. We have seen no changes in the dataset after day 7. Additionally, in the MAMS trial as all groups received the combination therapy from the first day, however there was still an agreement between the estimates based on the EBA trial and the MAMS trial. Therefore, the inclusion of the other TB drugs used for drug susceptible TB should not affect our conclusions substantially, however this cannot be ruled out. Furthermore, the inclusion of the other drugs makes it difficult to assess whether our results are true for rifampicin monotherapy by itself or just for combination therapy. Third, the PK PD model of heteroresistance currently cannot be used to predict TSCC. This is because in the model there are some subpopulations with very low susceptibility (16x MIC compared to the majority of the population) that are killed extremely slowly by antibiotics. These differences between the model and measurements can be due to the following: (i) the real susceptibility distribution of bacteria may be different, and subpopulations with very low susceptibility may not exist. (ii) In the model, the less susceptible subpopulations are modelled as having decreased effective antibiotic concentrations [52]. (iii) Immune system mediated killing of bacteria is not modelled as the killing rates for it are unknown. However, there is evidence that the immune system also plays a role in controlling TB infections [53–56]. Omitting these from the PK PD models can affect the estimated TSCC in various ways and may correct for an overestimated TSCC when accounting for subpopulation with >16MIC. This would imply that the immune system by itself may be able to handle very small bacterial populations.

Taken together, our results suggest that the bactericidal activity of TB treatments can be enhanced by higher rifampicin doses as well as by optimizing for treatment strategies and rifampicin formulations that allow peak concentrations above the MIC of all subpopulations of bacteria. This in turn may increase treatment success rates by reducing the time to sputum culture conversion: the time until no more bacteria are detected in the sputum. A shorter time to sputum conversion has been shown to be mildly correlated with treatment success [5]. This is supported by previous works on the same trial, which linked higher rifampicin exposures to an increased probability of earlier culture conversion [21], and increased time to positivity in liquid cultures [20]. Other studies have also shown that higher doses of rifampicin shorten treatment durations in mouse models of both *Mycobacterium tuberculosis* [22] and *Mycobacterium ulcerans* [57]. Finally, with a limited number of patients, in a different study we linked higher rifampicin doses per bodyweight to the rate of late killing [13]. Therefore, our work adds to the growing body of literature supporting optimized rifampicin doses for TB therapy and give hope that optimized higher doses could allow shortening treatment as is currently investigated in ongoing clinical trials (NCT02581527, NCT02410772, [24,47,57]).

## Acknowledgements

The original studies are part of the PanACEA consortium, which is part of the European and Developing Countries Clinical Trials Partnership programme (EDCTP) 1 (project codes IP.2007.32011.011, IP.2007.32011.012, and IP.2007.32011.013) and the EDCTP2 programme supported by the European Union (grant number TRIA2015-1102-PanACEA).

Antal Martinecz is currently an employee of AstraZeneca, however the work has been carried out prior to his employment.

We thank Christopher Fröhlich, Santiago Ramon Garcia, and Jingyi Liang, Sören Abel, Helen Stagg, Helen Jenkins, and Bill Hanage for their comments and discussions on the manuscript.

## Supplementary Material

**Figure S 1.**
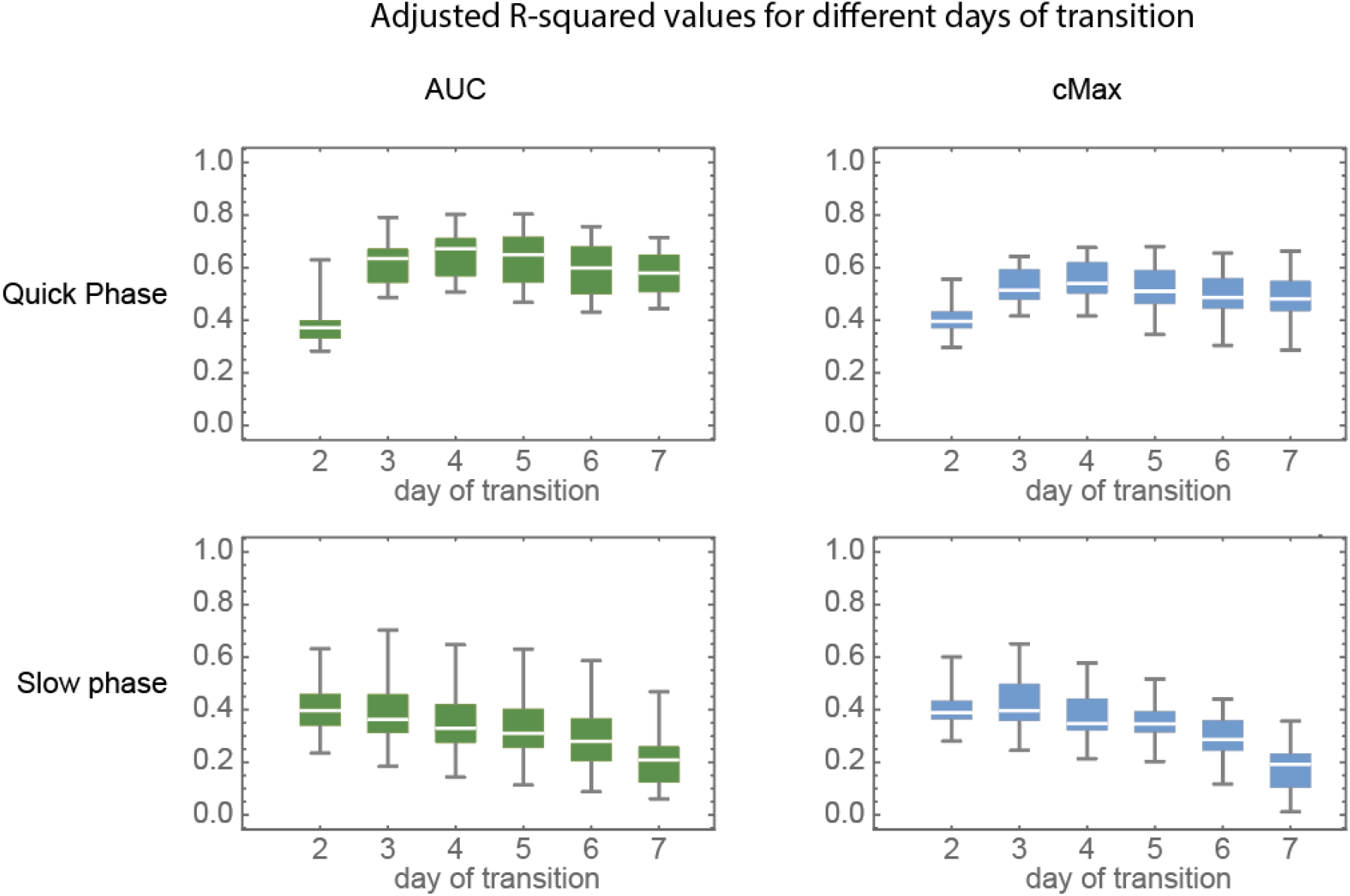
Adjusted R squared values for the fits on the quick and slow phases’ dependence on PK values. The different box-whisker plots in each figure correspond to a different set day of transition. These plots both show that values are consistently better predictors for the slope of the slow phase (based on adjusted R-squared values), as well as that we achieve the best fits for days 3 and 4.

**Figure S 2.**
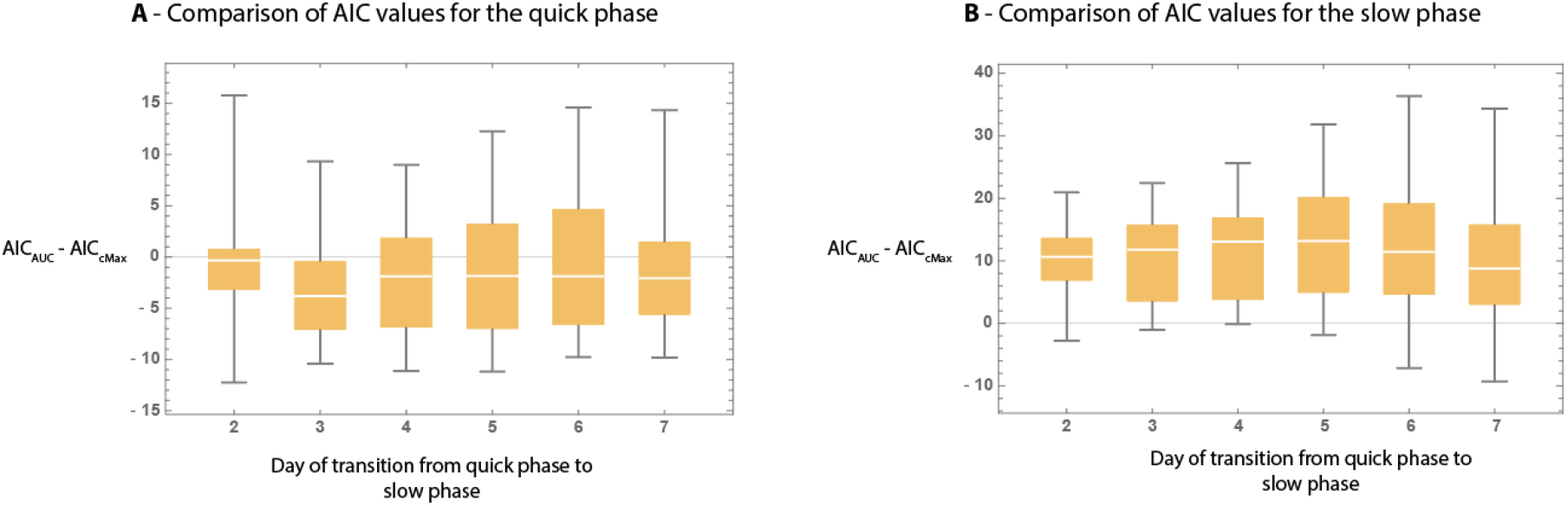
Difference of corrected AIC values for the fits on the slow and quick phases for different days of transition. Here, the positive values indicate that C_max_ is a better predictor, while negative values would indicate that AUC is a better predictor for the given phase.

**Figure S 3.**
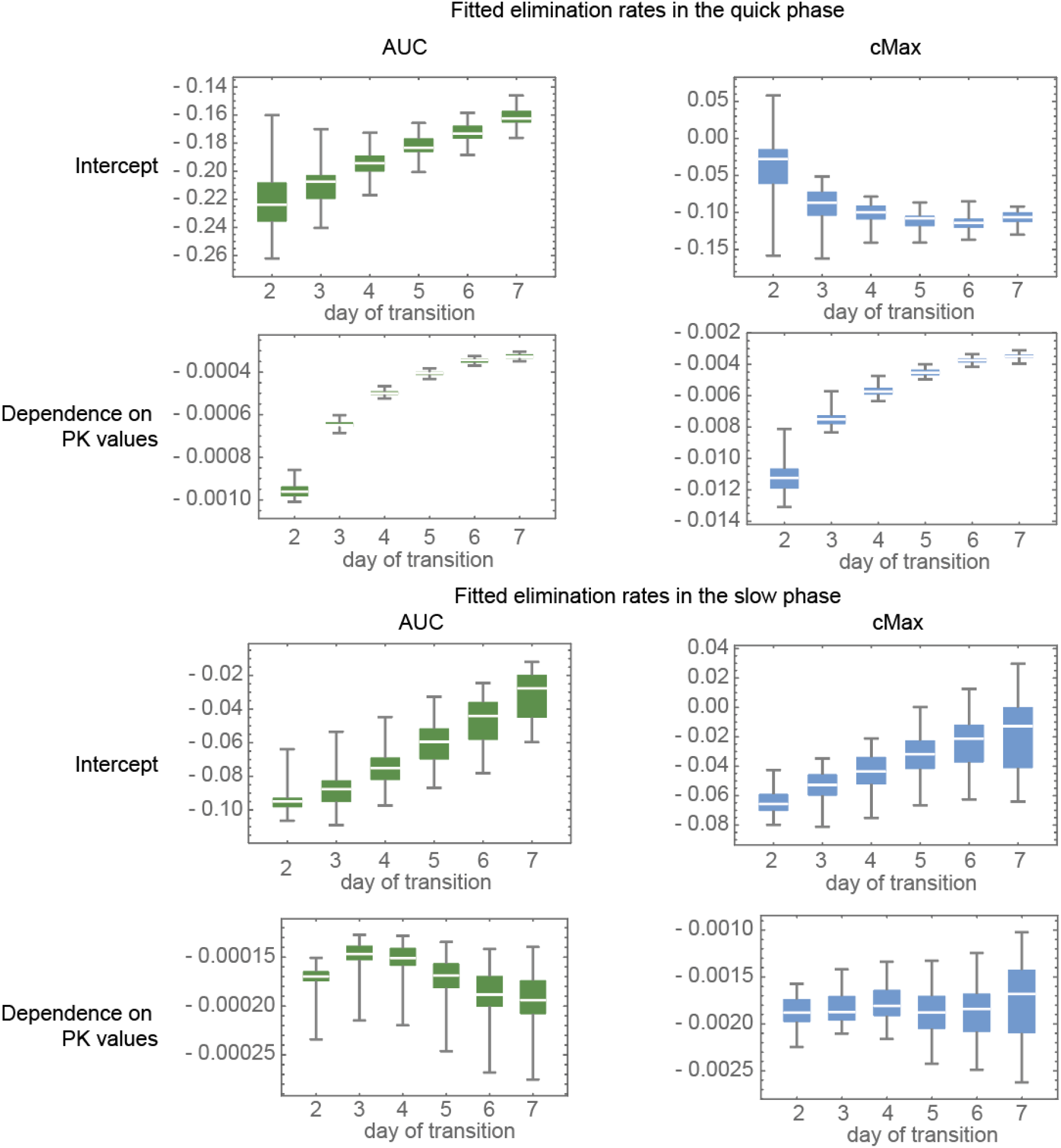
The determined fits of the biphasic curves for different days of transition (days 2-7, Y-axis) from quick to slow phase. The box-whisker plots represent the results from groupings from dividing the range of PK values into 10 – 40 equal intervals.

**Figure S 4.**
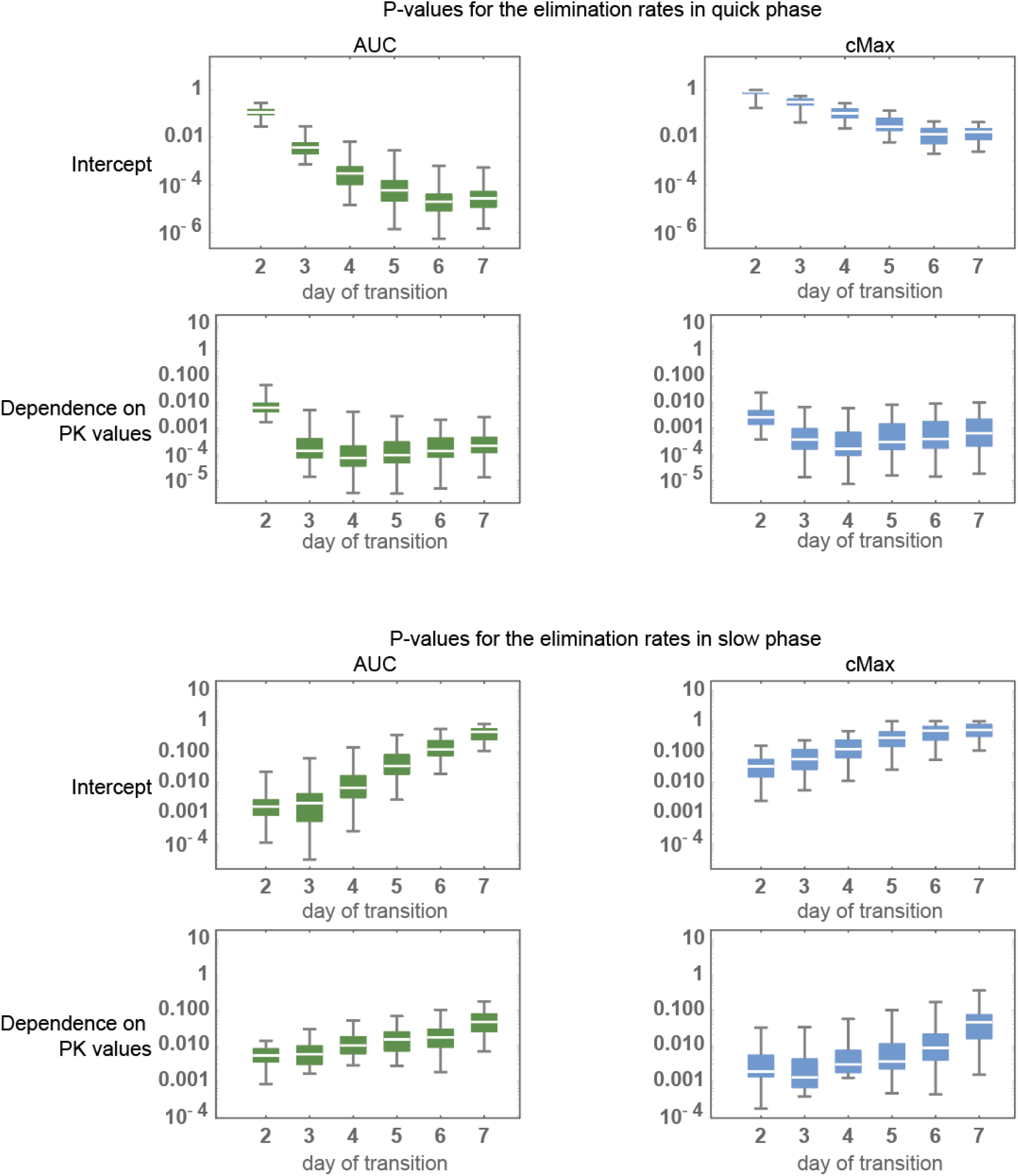
P-values for the fitted data with different day of transition (days 2-7, Y-axis) from quick to slow phase. The box-whisker plots represent the results from groupings from dividing the range of PK values into 10 – 40 equal intervals. These plots show that if we get the best fits if the days of transition are set to day 3 or 4. Furthermore these also show that C_max_ is consistently a better predictor for the slope of the slow phase than the AUC.

**Figure S 5.**
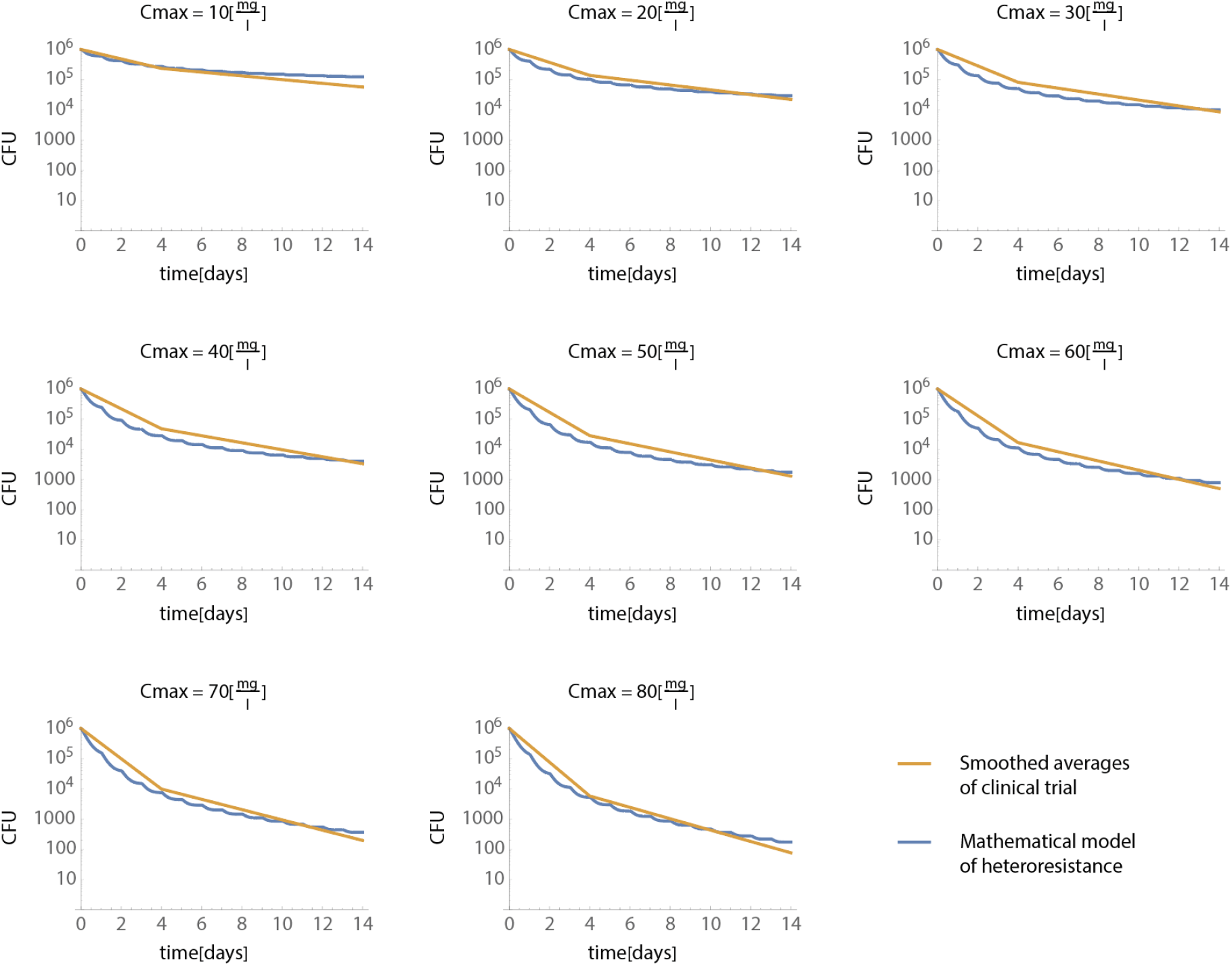
Side by side comparison of the smoothed averages of the clinical trial dataset and the mathematical model of heteroresistance. All figures show the observed/predicted bacterial counts (Y-axis) over time (X-axis). Each figure shows the same at different Cmaxes. The C_max_-es were chosen to be at regular intervals within the range of the clinical trial dataset (10-80 mg/l), the inputs doses for the mathematical models were chosen to achieve the same Cmaxes within the model.

**Figure S 6.**
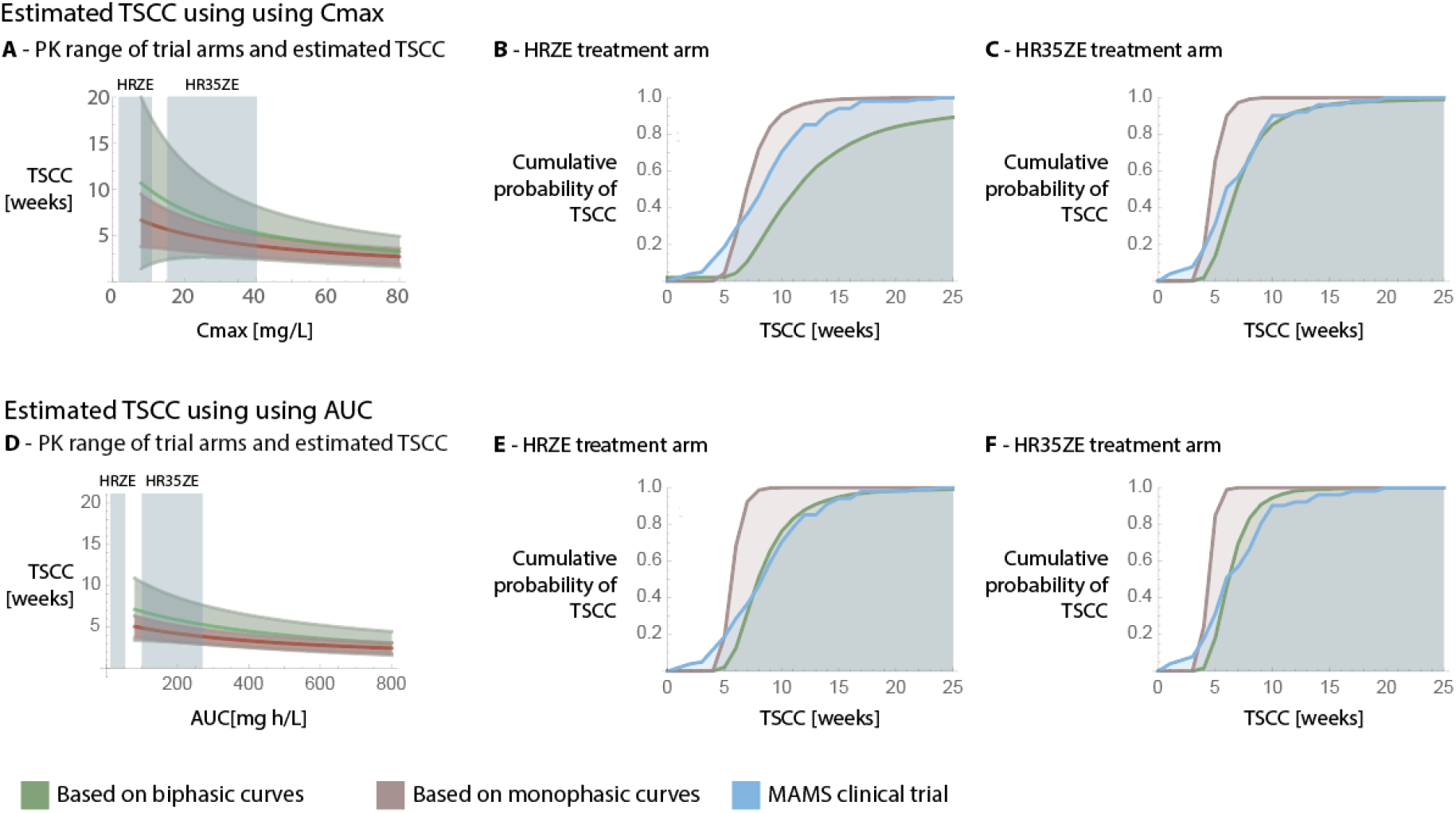
Estimated TSCC based on the statistical analysis, and comparison to measured TSCCs in a multi arm multistage (MAMS) clinical trial (NCT01785186, [25]). Note: this the left and right columns of this figure are the same as Figure 6. but extended with the standard dosing group for comparison. There, estimates relied on extrapolation outside the range of PK values of the EBA trials (see Figures A and D). On all of the plots the green curves show estimates based on biphasic curves (i.e taking the slowdown in decline into account), while red shows estimation based on monophasic curves (i.e. neglecting the possibility of a slowdown in decline). Blue always corresponds to data from the MAMS clinical trial. **Figures A** and **D** show the dependence of the predicted TSCC on the pharmacokinetic parameters (Cmax and AUC respectively), as well as the measured pharmacokinetic ranges for the HRZE and HR35ZE treatment arms in the MAMS clinical trial used for comparison (blue boxes). Here, the area around the estimates signify the 95% confidence interval around our estimates. The estimates are only shown within the parameter ranges of the EBA clinical trial. **Figures B**,**C**,**E**, and **F** show the cumulative probability (Y-axis) of TSCC (X-axis) based on the estimates within the PK ranges of the MAMS trial as well as the data from the MAMS trial itself. This is shown for both Cmax and AUC, as well as the HRZE, and HR35ZE treatment arms.

**Table S1.**
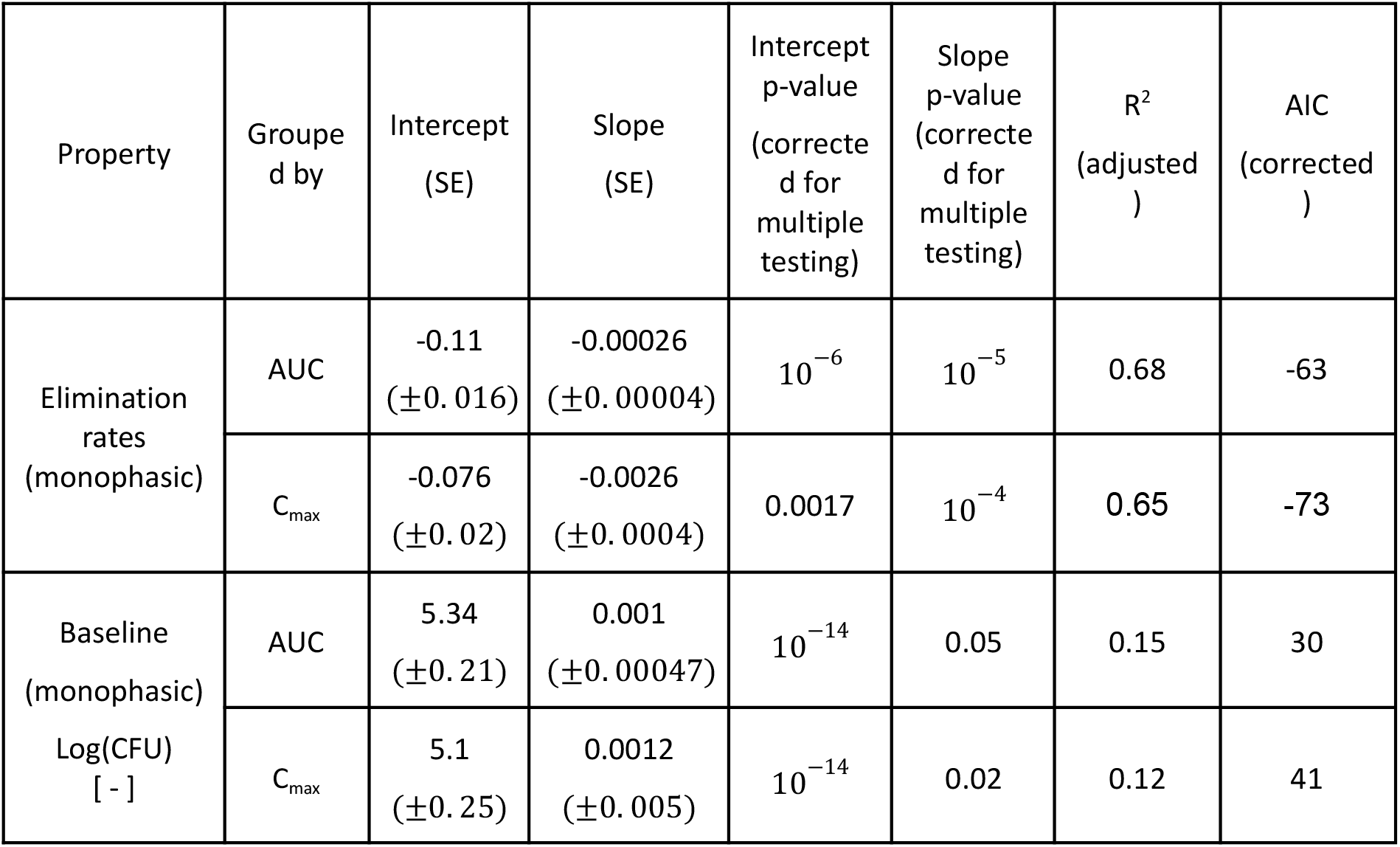
Summary of the median fitted values to the dataset across all groupings, when assuming a monophasic killing (fitting a straight line though the data). All the P values reported are corrected for multiple testing with the Benjamini-Hochberg method.

## References

1. Horsburgh CR, Barry CE, Lange C. Treatment of Tuberculosis. N Engl J Med 2015; 373:2149–2160.

2. Lawn SD, Zumla AI. Tuberculosis. The Lancet 2011; 378:57–72.

3. Dooley KE, Hanna D, Mave V, Eisenach K, Savic RM. Atdvancing the development of new tuberculosis treatment regimens: The essential role of translational and clinical pharmacology and microbiology. PLOS Med 2019; 16:e1002842.

4. Davies G, Boeree M, Hermann D, Hoelscher M. Accelerating the transition of new tuberculosis drug combinations from Phase II to Phase III trials: New technologies and innovative designs. PLOS Med 2019; 16:e1002851.

5. Horne DJ, Royce SE, Gooze L, et al. Sputum monitoring during tuberculosis treatment for predicting outcome: systematic review and meta-analysis. Lancet Infect Dis 2010; 10:387–94.

6. Shamputa IC, Jugheli L, Sadradze N, et al. Mixed infection and clonal representativeness of a single sputum sample in tuberculosis patients from a penitentiary hospital in Georgia. Respir Res 2006; 7. Available at: http://respiratory-research.biomedcentral.com/articles/10.1186/1465-9921-7-99. Accessed 7 April 2019.

7. Metcalfe JZ, Streicher E, Theron G, et al. Mycobacterium tuberculosis Subculture Results in Loss of Potentially Clinically Relevant Heteroresistance. Antimicrob Agents Chemother 2017; 61. Available at: http://aac.asm.org/lookup/doi/10.1128/AAC.00888-17. Accessed 7 April 2019.

8. Shockey AC, Dabney J, Pepperell CS. Effects of Host, Sample, and in vitro Culture on Genomic Diversity of Pathogenic Mycobacteria. Front Genet 2019; 10:477.

9. Balaban NQ, Helaine S, Lewis K, et al. Definitions and guidelines for research on antibiotic persistence. Nat Rev Microbiol 2019; Available at: http://www.nature.com/articles/s41579-019-0196-3. Accessed 18 April 2019.

10. Balaban NQ, Merrin J, Chait R, Kowalik L, Leibler S. Bacterial persistence as a phenotypic switch. Science 2004; 305:1622–1625.

11. Barr DA, Kamdolozi M, Nishihara Y, et al. Serial image analysis of Mycobacterium tuberculosis colony growth reveals a persistent subpopulation in sputum during treatment of pulmonary TB. Tuberculosis 2016; 98:110–115.

12. Saliba A-E, Li L, Westermann AJ, et al. Single-cell RNA-seq ties macrophage polarization to growth rate of intracellular Salmonella. Nat Microbiol 2016; 2:16206.

13. Abel zur Wiesch P, Abel S, Gkotzis S, et al. Classic reaction kinetics can explain complex patterns of antibiotic action. Sci Transl Med 2015; 7:287ra73.

14. Bergmiller T, Andersson AMC, Tomasek K, et al. Biased partitioning of the multidrug efflux pump AcrAB-TolC underlies long-lived phenotypic heterogeneity. Science 2017; 356:311–315.

15. Rego EH, Audette RE, Rubin EJ. Deletion of a mycobacterial divisome factor collapses single-cell phenotypic heterogeneity. Nature 2017; 546:153–157.

16. El-Halfawy OM, Valvano MA. Antimicrobial Heteroresistance: an Emerging Field in Need of Clarity. Clin Microbiol Rev 2015; 28:191–207.

17. Ley SD, de Vos M, Van Rie A, Warren RM. Deciphering Within-Host Microevolution of Mycobacterium tuberculosis through Whole-Genome Sequencing: the Phenotypic Impact and Way Forward. Microbiol Mol Biol Rev 2019; 83. Available at: http://mmbr.asm.org/lookup/doi/10.1128/MMBR.00062-18. Accessed 7 April 2019.

18. Sarathy J, Dartois V, Dick T, Gengenbacher M. Reduced Drug Uptake in Phenotypically Resistant Nutrient-Starved Nonreplicating Mycobacterium tuberculosis. Antimicrob Agents Chemother 2013; 57:1648–1653.

19. Magombedze G, Pasipanodya JG, Gumbo T. Bacterial load slopes represent biomarkers of tuberculosis therapy success, failure, and relapse. Commun Biol 2021; 4:664.

20. Svensson RJ, Svensson EM, Aarnoutse RE, et al. Greater Early Bactericidal Activity at Higher Rifampicin doses Revealed by Modeling and Clinical Trial Simulations. J Infect Dis 2018; :1–9.

21. Svensson EM, Svensson RJ, te Brake LHM, et al. The Potential for Treatment Shortening With Higher Rifampicin Doses: Relating Drug Exposure to Treatment Response in Patients With Pulmonary Tuberculosis. Clin Infect Dis 2018; 67.

22. Hu Y, Liu A, Ortega-Muro F, Alameda-Martin L, Mitchison D, Coates A. High-dose rifampicin kills persisters, shortens treatment duration, and reduces relapse rate in vitro and in vivo. Front Microbiol 2015; 6.

23. Martinecz A, Abel zur Wiesch P. Estimating treatment prolongation for persistent infections. Pathog Dis 2018; 76.

24. Boeree MJ, Diacon AH, Dawson R, et al. A dose-ranging trial to optimize the dose of rifampin in the treatment of tuberculosis. Am J Respir Crit Care Med 2015; 191:1058–1065.

25. Boeree MJ, Heinrich N, Aarnoutse R, et al. High-dose rifampicin, moxifloxacin, and SQ109 for treating tuberculosis: a multi-arm, multi-stage randomised controlled trial. Lancet Infect Dis 2016; 17:39–49.

26. Bowness R, Boeree MJ, Aarnoutse R, et al. The relationship between mycobacterium tuberculosis mgit time to positivity and cfu in sputum samples demonstrates changing bacterial phenotypes potentially reflecting the impact of chemotherapy on critical sub-populations. J Antimicrob Chemother 2015; 70:448–455.

27. Magombedze G, Pasipanodya JG, Srivastava S, et al. Transformation Morphisms and Time-to-Extinction Analysis That Map Therapy Duration From Preclinical Models to Patients With Tuberculosis: Translating From Apples to Oranges. Clin Infect Dis 2018; 67:S349–S358.

28. Chao MC, Rubin EJ. Letting Sleeping dos Lie: Does Dormancy Play a Role in Tuberculosis? Annu Rev Microbiol 2010; 64:293–311.

29. Cadosch D, Abel zur Wiesch P, Kouyos R, Bonhoeffer S. The Role of Adherence and Retreatment in De Novo Emergence of MDR-TB. PLoS Comput Biol 2016; 12:1–19.

30. de Steenwinkel JEM, de Knegt GJ, ten Kate MT, et al. Time-kill kinetics of anti-tuberculosis drugs, and emergence of resistance, in relation to metabolic activity of Mycobacterium tuberculosis. J Antimicrob Chemother 2010; 65:2582–2589.

31. Marcel N, Nahta A, Balganesh M. Evaluation of killing kinetics of anti-tuberculosis drugs on Mycobacterium tuberculosis using a bacteriophage-based assay. Chemotherapy 2008; 54:404–411.

32. Aljayyoussi G, Jenkins VA, Sharma R, et al. Pharmacokinetic-Pharmacodynamic modelling of intracellular Mycobacterium tuberculosis growth and kill rates is predictive of clinical treatment duration. Sci Rep 2017; 7:1–11.

33. Grosset J. Bacteriologic basis of short-course chemotherapy for tuberculosis. Clin Chest Med 1980; 1:231–241.

34. Koul A, Vranckx L, Dendouga N, et al. Diarylquinolines Are Bactericidal for Dormant Mycobacteria as a Result of Disturbed ATP Homeostasis. J Biol Chem 2008; 283:25273–25280.

35. Gengenbacher M, Rao SPS, Pethe K, Dick T. Nutrient-starved, non-replicating Mycobacterium tuberculosis requires respiration, ATP synthase and isocitrate lyase for maintenance of ATP homeostasis and viability. Microbiology 2010; 156:81–87.

36. Strydom N, Gupta SV, Fox WS, et al. Tuberculosis drugs’ distribution and emergence of resistance in patient’s lung lesions: A mechanistic model and tool for regimen and dose optimization. PLOS Med 2019; 16:e1002773.

37. Gumbo T, Louie A, Deziel MR, et al. Concentration-Dependent Mycobacterium tuberculosis Killing and Prevention of Resistance by Rifampin. Antimicrob Agents Chemother 2007; 51:3781–3788.

38. Andersson DI, Nicoloff H, Hjort K. Mechanisms and clinical relevance of bacterial heteroresistance. Nat Rev Microbiol 2019; Available at: http://www.nature.com/articles/s41579-019-0218-1. Accessed 1 July 2019.

39. Pu Y, Zhao Z, Li Y, et al. Enhanced Efflux Activity Facilitates Drug Tolerance in Dormant Bacterial Cells. Mol Cell 2016; 62:284–294.

40. Band VI, Weiss DS. Heteroresistance: A cause of unexplained antibiotic treatment failure? PLOS Pathog 2019; 15:e1007726.

41. Lenaerts A, Barry CE, Dartois V. Heterogeneity in tuberculosis pathology, microenvironments and therapeutic responses. Immunol Rev 2015; 264:288–307.

42. Operario DJ, Koeppel AF, Turner SD, et al. Prevalence and extent of heteroresistance by next generation sequencing of multidrug-resistant tuberculosis. PLOS ONE 2017; 12:e0176522.

43. Mandal S, Njikan S, Kumar A, Early JV, Parish T. The relevance of persisters in tuberculosis drug discovery. Microbiology 2019; 165:492–499.

44. Genestet C, Hodille E, Barbry A, et al. Rifampicin exposure reveals within-host Mycobacterium tuberculosis diversity in patients with delayed culture conversion. PLOS Pathog 2021; 17:e1009643.

45. Dartois V. The path of anti-tuberculosis drugs: from blood to lesions to mycobacterial cells. Nat Rev Microbiol 2014; 12:159–167.

46. Prideaux B, Via LE, Zimmerman MD, et al. The association between sterilizing activity and drug distribution into tuberculosis lesions. Nat Med 2015; 21:1223–1227.

47. Dorman SE, Nahid P, Kurbatova EV, et al. High-dose rifapentine with or without moxifloxacin for shortening treatment of pulmonary tuberculosis: Study protocol for TBTC study 31/ACTG A5349 phase 3 clinical trial. Contemp Clin Trials 2020; 90:105938.

48. Imperial MZ, Nahid P, Phillips PPJ, et al. A patient-level pooled analysis of treatment-shortening regimens for drug-susceptible pulmonary tuberculosis. Nat Med 2018; 24:1708–1715.

49. Diacon AH, Dawson R, Von Groote-Bidlingmaier F, et al. Randomized dose-ranging study of the 14-day early bactericidal activity of bedaquiline (TMC207) in patients with sputum microscopy smear-positive pulmonary tuberculosis. Antimicrob Agents Chemother 2013; 57:2199–2203.

50. Diacon AH, van der Merwe L, Demers A-M, Von Groote-Bidlingmaier F, Venter A, Donald PR. Pre-treatment mycobacterial sputum load influences individual on-treatment measurements. Tuberc Edinb Scotl 2014; 94:690–694.

51. De Jager V, van der Merwe L, Venter A, Donald PR, Diacon AH. Time Trends in Sputum Mycobacterial Load and Two-Day Bactericidal Activity of Isoniazid-Containing Antituberculosis Therapies. Antimicrob Agents Chemother 2017; 61.

52. Martinecz A, Clarelli F, Abel S, Abel zur Wiesch P. Reaction Kinetic Models of Antibiotic Heteroresistance. Int J Mol Sci 2019; 20:3965.

53. Torrelles JB, Schlesinger LS. Integrating Lung Physiology, Immunology, and Tuberculosis. Trends Microbiol 2017; 25:688–697.

54. Furin J, Cox H, Pai M. Tuberculosis. The Lancet 2019; 393:1642–1656.

55. the Catalysis TB–Biomarker Consortium, Malherbe ST, Shenai S, et al. Persisting positron emission tomography lesion activity and Mycobacterium tuberculosis mRNA after tuberculosis cure. Nat Med 2016; 22:1094–1100.

56. Lin PL, Ford CB, Coleman MT, et al. Sterilization of granulomas is common in active and latent tuberculosis despite within-host variability in bacterial killing. Nat Med 2014; 20:75–79.

57. Omansen TF, Almeida D, Converse PJ, et al. High-Dose Rifamycins Enable Shorter Oral Treatment in a Murine Model of Mycobacterium ulcerans Disease. Antimicrob Agents Chemother 2018; 63:e01478–18.

